# GNAQ Induces Melanomagenesis in Mitfa-Independent Melanocyte Progenitors in a Zebrafish Model of Uveal Melanoma

**DOI:** 10.1101/2025.05.05.652300

**Authors:** Julius I. Yevdash, Delaney Robinson, Rachel Moore, Zhijie Li, Katelyn R. Campbell-Hanson, Danielle Gutelius, Stephen P. G. Moore, Dylan Friend, Isaac O’Toole, Brynnon E Harman, Collin Montgomery, Robert A. Cornell, Alexander Birbrair, Jesse D. Riordan, Adam J. Dupuy, Elaine M. Binkley, Robert F. Mullins, Deborah Lang, Ronald J. Weigel, Colin Kenny

**Author notes:** The authors declare no potential conflicts of interest.

## Abstract

Melanocytes reside in diverse microenvironments that influence their susceptibility to oncogenic transformation; however, investigation of rare melanoma subsets has been limited by the lack of suitable pre-clinical animal models. Here, we developed a primary, immune-competent zebrafish model to study uveal melanoma (UM) using choroidal melanocyte-targeted injection and electroporation of plasmids encoding human GNAQ^Q209L^ together with CRISPR/Cas9 cassettes for somatic tumor suppressor gene deletion. Single-cell transcriptional profiling of primary melanocytes and melanoma derived from the eye and skin revealed distinct transcriptional programs, with epithelial-to-mesenchymal transition pathways enriched in ocular tumors. In addition, choroidal fibroblasts from tumor-bearing eyes exhibited marked transcriptional changes, including increased fibronectin and collagen expression, consistent with stromal remodeling. Given prior associations between *mitfa* loss and accelerated GNAQ^Q209L^ tumor onset, the model was applied to determine whether melanocyte differentiation state contributes to the emergence of GNAQ-driven tumors. The increased susceptibility resulted from expansion of Mitfa-independent melanocyte progenitor populations in germline *mitfa* mutant zebrafish, rather than somatic *mitfa* loss in differentiated melanocytes, as conditional, melanocyte-specific *mitfa* deletion in adult zebrafish did not accelerate tumor growth. Furthermore, *pax3a*-positive melanocyte progenitor cells in *mitfa*-deficient zebrafish embryos and adult eyes and skin were highly susceptible to transformation induced by GNAQ^Q209L^ but not BRAF^V600E^. Analogous *PAX3* positive populations were also identified in mouse and human single-cell transcriptomic datasets. Collectively, these findings establish a critical role for Mitfa-independent melanocyte progenitors in UM pathogenesis.

**Statement of Significance:** Choroid-targeted GNAQ^Q209L^ expression induces anatomically correct uveal melanoma in adult zebrafish, with germline *mitfa* deletion expanding mitfa-independent melanocyte progenitors with enhanced susceptibility that are transcriptionally distinct from the subpopulation transformed by BRAF^V600E^.

## Introduction

Uveal melanoma (UM) is a rare cancer of the melanocytes in the uveal tract. The majority of UM’s (∼90%) occur in the choroid, the vascular structure supplying the outer retina, with the remainder arising from the ciliary body or the iris.^1^ Oncogenic mutations in *GNAQ* or its paralog *GNA11* are observed in up to 90% of UMs but are rare in cutaneous melanoma (CM).^2^ Such mutations reduce GTPase activity of the heterotrimeric G protein alpha subunit, resulting in constitutive activation that drives cellular proliferation through downstream signaling pathways, including MAPK, YAP/TAZ, and PI3K.^3,4^ The transcriptional mechanisms of UM onset and progression remain understudied, in part, because existing preclinical models inadequately capture the earliest events of ocular melanocyte transformation within the correct microenvironment.

A recent study by Phelps et al.^5^ showed that loss of *mitfa,* which is considered the master regulator of the melanocyte lineage, accelerates tumorigenesis in transgenic *Tg(mitfa-GNAQ^Q209L^); mitfa^-/-^; tp53^-/-^* zebrafish compared to *Tg(mitfa-GNAQ^Q209L^); tp53^-/-^* controls. These findings indicate that, in the context of tumor-suppressor loss, oncogenic GNAQ can drive tumor formation independently of differentiated melanocytes. This contrasts with *BRAF^V600E^*-induced melanoma, where Mitfa is required for tumor formation.^6^ Such observations imply that melanocyte progenitor cells, which persist in *mitfa*-mutant zebrafish, may serve as the cells of origin for GNAQ-induced transformation. While transgenic models have been incredibly informative for studying spontaneous tumor formation, germline loss of *mitfa* or tumor suppressor genes, together with melanocyte-lineage expression of oncogenes throughout development, may unintentionally perturb melanocyte lineage specification. Consequently, these models may capture tumorigenesis in a lineage already altered during development rather than melanoma initiated by somatic mutations in normal melanocytes. Conditional knockout approaches as well as lineage restricted expression of oncogenes in the appropriate microenvironment will improve our understanding of UM pathogenesis.

Zebrafish are a well-established model for studying melanocytes and melanoma due to conserved melanocyte differentiation pathways, efficient genetic manipulation, and the availability of transparent lines that facilitate high-resolution live-cell imaging. Zebrafish have three types of pigment cells; dark melanocytes, iridescent iridophores, and yellow xanthophores, that arise from multipotent, neural crest-derived pigment progenitors.^7^ To study early events in UM, we developed a novel zebrafish model that utilizes transgene electroporation of adult zebrafish (TEAZ)^8^ to express the human *GNAQ^Q209L^* oncogene and conditionally delete tumor suppressor genes in cells of the melanocyte lineage within the skin and eye. Choroidal targeted injection and electroporation drive primary UM onset from the same anatomical site as human UM. In this study, we demonstrate the existence of a melanocyte progenitor population in zebrafish eyes with increased sensitivity to transformation by GNAQ^Q209L^. Such progenitor cells are expanded in *mitfa* deficient zebrafish and result in reduced GNAQ^Q209L^-induced tumor latency compared to wild-type animals.

## Materials and Methods

Plasmid constructs and zebrafish generated in this study may be obtained by a request to the lead contact.

### Data and code availability

Datasets generated in this article are available from NCBI GEO accession number GEO: GSE307227. This study does not report original code. Any additional information required to reanalyze the data reported in this work paper is available from the lead contact upon request.

### Human donor eye procurement and histology

Human donor eyes were obtained from the Iowa Lions Eye Bank with full informed consent from the next of kin for use in research. All procedures adhered to the tenets of the Declaration of Helsinki and institutional guidelines. Globes were processed for histological analysis in the laboratory of Dr. Mullins at the University of Iowa. Tissues were fixed, paraffin embedded, sectioned, and stained with hematoxylin and eosin (H&E) using standard protocols.

### Zebrafish husbandry

All zebrafish used in this study were bred and maintained at the University of Iowa Animal Care Facility. The zebrafish are kept consistently at 28°C, 7.4 pH, and controlled salt concentrations and cyclically receive 14 h of light followed by 10 hours of darkness. All zebrafish experiments were performed in compliance with the ethical regulations of the Institutional Animal Care and Use Committee at the University of Iowa and in compliance with NIH guidelines (protocol #3022523). Zebrafish embryos were maintained at 28.5°C and staged by hours or days post-fertilization (hpf or dpf).

### Zebrafish mutant lines

Wild-type zebrafish in this study include the *AB* line (RRID:ZFIN_ZDBGENO-960809-7) and the *WIK* line (RRID:ZIRC_ZL84). Transgenic lines include *casper* (*mitfa*^w2/w2^; *mpv17*^-/-^)^9,10^ (RRID:ZIRC_ZL1714), *nacre* (*mitfa*^w2/w2^)^11^ (RRID:ZIRC_ZL2104), *Tg*(*mitfa*:GFP)^18^, and *Tg*(*mitfa*:GFP); *mitfa*^w2/w2^. TEAZ-Eye and TEAZ-Skin was performed on adult zebrafish (inclusion criteria: greater than 4 months postfertilization and less than 1.5 years old). Similar proportions of male and female zebrafish were used within each experiment.

### Induction of melanoma using Strategies A and B

To model GNAQ-driven melanoma, we generated *mitfa* promoter–driven plasmid systems expressing human oncogenic GNAQ^Q209L^, CRISPR/Cas9, and guide RNAs (gRNAs) targeting *tp53, ptena, and ptenb*. Two delivery strategies were used. In Strategy A, a single plasmid contained two expression cassettes under the mitfa promoter: one encoding GNAQ^Q209L^ fused to a T2A self-cleaving peptide followed by eGFP, and a second encoding Cas9, together with three U6-driven gRNAs targeting *tp53* and *ptena/b*. Because T2A-mediated cleavage leaves residual C-terminal amino acids that may alter GNAQ oncogenic activity, we also employed an alternative multi-plasmid approach (Strategy B). This system consisted of five separate plasmids: mitfa:*GNAQ^Q209L^*, mitfa:GFP, mitfa:Cas9, U6:gRNA-*tp53*, U6:gRNA-*ptena*, and U6:gRNA-*ptenb*. Plasmids expressing mitfa:GNAQ^Q209L^ and mitfa:GFP were generously provided by Dr. Jacqueline Lees (MIT), and plasmids encoding Cas9 and gRNAs were provided by Dr. Richard White (University of Oxford). Strategy B was designed to avoid unintended C-terminal modification of GNAQ and to improve electroporation efficiency due to reduced plasmid size. Fish were monitored weekly for tumor formation at the electroporation site. Tumor onset was defined as the first appearance of a visible, expanding pigmented or translucent mass. Tumor latency was calculated as days post-electroporation to first detection. Kaplan–Meier analyses were used to compare tumor onset between experimental groups.

### Cloning

The MiniCoopR^6^ vector (MiniCoopR 2x*U6*:gRNA, *mitfa*:Cas9 was a gift from Leonard Zon Addgene plasmid # 118844) was modified to replace the *mitfa* coding sequence with *GNAQ*^Q209L^ by removing *mitfa* using StuI and SmaI restriction enzymes (NEB). Gibson assembly was used to insert *GNAQ*^Q209L^:T2A:GFP DNA as a gBlock hifi (IDT) with sequence on the 3’ (GAAGCTAACACATAGTTGAAC) and 5’ (CAATGCCAACTAAATTTCATG) as overhangs.

Gibson assemblies were transformed using NEB 5-alpha competent *E. coli* cells via heat shock. *GNAQ*^Q209L^:T2A:GFP insertion was confirmed using Sanger sequencing (Primers: CAAGGAAGCCCGGCGGATCAACGA, CATACTTGTATGGGATCTTGAGTGTGTCCA, GTGGAGTCAGACAATGAGAACCGAATGGAG, CAAGCTGGAGTACAACTACAA).

*mitfa* sgRNA was designed using IDT’s CRISPR-Cas9 guide RNA design tool (sequence: ATGGACAAAGCTGGACCATG). *U6*:sgRNA-*ptena* was linearized by PCR (forward primer: GTTTAAGAGCTATGCTGGAAACAGCAT, reverse primer: GAACAAAGAGCTGGAGGGAG). Linear plasmid was purified from a 2% agarose gel using the Monarch GNA Gel Extraction Kit (#T1020S). ssDNA sequences for sgRNA-*mitfa* were annealed together in NE buffer r2.1. Gibson assembly was used to insert sgRNA-*mitfa* into linearized U6 plasmid. Assembled product was transformed into NEB 5-alpha competent *E. coli* cells via heat shock and selected with 50 ug/mL streptomycin. Insertion of sgRNA-*mitfa* was confirmed using PCR amplification of lysed bacteria from single colonies (forward primer: ATGGACAAAGCTGGACCATG, reverse primer: CTTAGCTGGATAACGCCAC). Positive products were isolated using the QIAGEN Plasmid Maxi Kit. Entire plasmid was sequenced using Plasmidsaurus sequencing service.

### Transgene Electroporation of Adult Zebrafish Skin (TEAZ-Skin)

Adult zebrafish were anesthetized in 0.2% Tricaine prior to upright placement on a prewetted Kimwipe. Each injection consisted of 1000 ng total DNA: 100 ng of Tol2 plasmid and an equal division of the remaining 900 ng for the remaining constructs. Injection stocks contained a small volume (<1uL, 1:50 dilution) of green food coloring to visualize localization of injection. Injection volumes ranged from 1 to 1.5 uL depending on the injection mix, keeping 1000 ng total constant. Needles were pulled from 10 cm long filamented borosilicate glass capillaries at a pressure of 500, heat of 650, pull strength of 100, velocity of 200, and time delay of 40 in a Model P-97 Flaming/Brown micropipette puller from Sutter Instrument Co. Following injection, zebrafish were immediately electroporated (<1 min) by placing the cathode probe of ultrasound gel-coated Platnium Tweezertrode, 3MM, on the side of the injection site and the anode probe on the other side of the fish body. Electroporation was achieved with 5 60-ms pulses of 40 volts with a 1s pulse interval from a ECM 830 Electro Square Porator from BTX Harvard Apparatus. Following electroporation, zebrafish are placed into a recovery tank with fresh, gently stirred system water and are monitored until normal swimming resumes. Electroporated zebrafish are imaged weekly. *mitfa*:*GNAQ*^Q209L^; *mitfa*:GFP plasmid constructs were kindly provided by Dr. Jacqueline A. Lees (Massachusetts Institute of Technology, Cambridge, USA). *mitfa*:Cas9, *mitfa:*tdTomato, PCS2FA Tol2, *U6*:gRNA-*tp53, U6*:gRNA-*ptena,* and *U6*:gRNA-*ptenb* plasmids were kindly provided by Dr. Richard White (Sloan Kettering Institute, New York, USA).

### Transgene Electroporation of Adult Zebrafish Eye (TEAZ-Eye)

Pre-injection preparation for eye injections is the same as for skin injections including: 0.2% Tricaine anesthesia, needle size, and injection concentrations. To inject the eye, a blunt spherical head of a pin was gently pressed against the bottom half of the zebrafish eyeball to reveal the dorsal posterior regions of the eye. The needle was gently pressed into this location, where the cornea is softer, just through the sclera, keeping the needle as superficial as possible as to not damage the back of the eye. Injection volumes were around 0.25 uL and were terminated with a slight visual swelling of the eye. Following injection zebrafish were immediately electroporated (<1 min) by the cathode probe of ultrasound gel-coated Platnium Tweezertrode, 1MM, on the superior side of the eye nearest the injection and the anode probe on the inferior side of the eye. Electroporation was achieved with 3 50-ms pulses of 75 volts with a 1s pulse interval from a ECM 830 Electro Square Porator from BTX Harvard Apparatus. Recovery and imaging procedures for these fish did not differ from the skin injections.

### Imaging and image processing

Injected zebrafish were imaged using an upright Leica M205 FCA microscope with brightfield and GFP filters. Zebrafish were briefly anesthetized with 0.2% Tricaine and positioned on a Kimwipe mound for upright orientation. Images were taken and processed through the LAS X software. All histology slides were imaged using brightfield, DAPI, and GFP filters on an EVOS M5000 microscope from Invitrogen for individual images or on a Leica DMi8 microscope and stitched together by LAS X Thunder Imaging software for full tissue images.

### H&E and IF – zebrafish tissue

Whole zebrafish or zebrafish eyes were fixed in 4% paraformaldehyde for 48 hours at 4°C and paraffin embedded in the University of Iowa Pathology Core. Fish were sectioned at 7 µM thickness and placed on Superfrost Plus Microscope slides from Fisher Scientific, baked at 60°C, stained with hematoxylin and eosin, and mounted with Cytoseal 60 from Epredia. For immunofluorescence, sections were deparaffinized and hydrated followed by heat-induced antigen retrieval with 10 mM Na-citrate buffer and blocking with 3% milk in 0.1% Tween 20. slides were stained with primary antibodies against GFP (NB600-308 Rabbit anti-GFP polyclonal from Novus Biologicals, RRID:AB_10003058, at 1:250) overnight at 4°C and then fluorescent secondary antibodies against rabbit IgG (Alexa Fluor 488 goat anti-rabbit IgG, RRID:AB_2576217, at 1:500 from a 2 mg/mL stock) for 1 hr at room temperature. RPE autofluorescence was minimized using the Vector TrueVIEW Autofluorescence Quenching Kit. Sections were counterstained with DAPI, mounted using VECTASHIELD Vibrance Antifade Mounting Medium, and imaged the same day.

### IF - mouse eye tissue

C57BL/6 mice (RRID:MGI:2159769) eyes were collected, formalin-fixed, paraffin-embedded, and sectioned into 7 µm-thick sections. Sections were deparaffinized and hydrated followed by heat-induced antigen retrieval with citrate buffer (Vector Laboratories, Inc.), permeabilized with 1% horse serum in 0.2% Triton-X, blocked with 5% Horse serum in 0.2% Triton-X, and incubated with PAX3 primary antibody (Invitrogen 38-1801 Rabbit anti-PAX3 polyclonal, RRID:AB_2533359, at 1:200) overnight at 4°C. Samples were incubated with DyLightTM 488-labeled secondary antibody (Vector Laboratories, DI-1088-1.5, horse anti-rabbit, RRID:AB_2336403, at 1:2000) for 1 hr at room temperature, treated with autofluorescence quenching kit (Vector Laboratories, Inc. SP-8400-15), and cover slips fitted using Vectashield Antifade Mounting Medium with DAPI (Vector Laboratories, Inc. H-1000-10).

### Kaplan-Meier analysis

Zebrafish were followed for up to 70 weeks and tumor-free survival was analyzed with the Kaplan-Meier method. Tumors were defined as the first presence of a GFP-positive nodular mass at the injection site. Zebrafish were excluded from curves if they perished or were euthanized without any visible signs of tumors during the study. Differences between injection groups were analyzed using log-rank statistics.

### Single cell suspension and single-cell RNA Seq analysis

Zebrafish were euthanized with 0.4% Tricaine for 10 mins followed by immersion in ice water until no opercular movement was observed for at least 30 min as detailed in our IACUC protocol. Tumors were harvested and minced into small pieces. Minced tumors were digested for 1 hour at 37°C with collagenase and hyaluronidase in HBSS with 2% FBS (HF). Suspensions were washed with 1:4 HF:NH_4_Cl and then further digested for 5 min with pre-warmed trypsin. Following trypsin inactivation, cell suspensions were washed twice with 10% FBS in HBSS.

Cellular suspensions were loaded on a 10x Genomics Chromium instrument to generate single-cell gel beads in emulsion (GEMs). Approximately 10,000-20,000 cells were loaded per channel depending on the 10x kit used. See figures S5 and S12 for kit information per sample. Next GEM kit: targeted cell number = 10,000. Single-cell RNA-Seq libraries were prepared using Single Cell 3′ Reagent Kits v2: Chromium Single Cell 3′ Library & Gel Bead Kit v2, PN-120237; Single Cell 3′ Chip Kit v2, PN-120236; and i7 Multiplex Kit, PN-120262 (10x Genomics) and following the Single Cell 3′ Reagent Kits v2 User Guide (Manual Part # CG00052 Rev A). GEM-X kit: targeted cell number = 20,000. Single-cell RNA-Seq libraries were prepared using Chromium GEM-X Single Cell 3′ Reagent Kits v4: Library Construction Kit C, PN-1000694; Single Cell 3’ GEM Kit v4, PN-1000693; Single Cell 3’ Gel Bead Kit v4, PN-2001128; and Dual Index Kit TT Set A, PN-3000431; and following the Chromium GEM-X Single Cell 3′ Reagent Kits v4 User Guide (Manual Part # CG000731 Rev B). Libraries were sequenced on an Illumina HiSeq 4000 as 2L×L150 paired-end reads. Sequencing results were demultiplexed and converted to FASTQ format using Illumina bcl2fastq software.

A custom reference genome for the Zebrafish was constructed with GRCz11 primary assembly (Ensembl, RRID:SCR_002344) using Cell Ranger (Cell Ranger mkref function). 10X Genomics scRNA-Seq reads were then processed and aligned to this reference with Cell Ranger (Cell Ranger Count function). Further analysis and visualization was performed using Seurat (v4.1.0)^12^, cells with fewer than 200 RNA feature counts and greater than 5% mitochondrial contamination were removed with filtering (greater than 15% for wildtype eye samples). RNA counts were normalized, and FindVariableFeatures was run with the following parameters: selection.method = “vst”, nfeatures = 2,000. The cells were originally clustered in a UMAP by RunPCA then RunUMAP with dims 1:30. FindNeighbors was run with dims 1:30 followed by FindClusters using a resolution of 0.4. These methods identified up to 20 clusters in individual samples.

For comparative analysis, RNA-seq datasets were integrated together using IntegrateData with dims 1:30. New clusters were generated as before. Clusters were annotated using FindAllMarkers and the top 25 enriched genes in each cluster were compared to the Daniocell dataset^13^ and Spectacle^14^. Cluster names were given according to the closest matched cellular population between these two datasets. To determine enriched pathways, *danio rerio* gene names were converted into human Ensembl gene codes with g:Profiler^15^ and then data were analyzed with the use of QIAGEN IPA (QIAGEN Inc)^16^ and GSEA.^17^ Pseudotime was performed using Monocle3 (v0.2.3.0).^18^

### Verification of sgRNAs

Zebrafish were euthanized with 0.4% Tricaine for 10 mins followed by immersion in ice water until no opercular movement was observed for at least 10 min as detailed in the IACUC protocol. Tumors were harvested and minced into small pieces. DNA was harvested according to the DNeasy Blood and Tissue Kit from Qiagen. Regions surrounding cut sites were PCR-amplified and the purity of the PCR amplicon was confirmed with agarose gel electrophoresis. Samples were cleaned up with the QIAquick PCR Purification Kit from Qiagen and sent for Sanger sequencing at the Iowa Institute of Human Genetics. Gene editing was confirmed with DECODR INDEL analysis by comparing to wildtype sequence^19^. PCR primer sequences follow: *mitfa* PCR F: CCGGTATGTATTCACATTGTCTTG *mitfa* PCR R: TCTTGCTTAGGATGCCTATGTATT *ptena* PCR F: CATCCCACCAAGTGAGGTTAAAC *ptena* PCR R: CACATACACAGTCAAGGGTGAG *ptenb* PCR F: CAGTTCTGTTGCACCCAATAAG *ptenb* PCR R: CTGGTGGTGTTGAGGCTATAAAG *tp53* PCR F: AAGTATTCAGCCCCCAGGTG *tp53* PCR R: CGCTTTTGACTCACAGTGCAAG

## Results

### Generation of GNAQ-driven melanoma in adult zebrafish

To model GNAQ-driven melanoma in adult zebrafish, we generated *mitfa* promoter-driven plasmid systems expressing oncogenic GNAQ^Q209L^, Cas9, and gRNAs targeting *tp53* and *ptena/b* (**Figure 1A**). We tested two delivery strategies: a single-plasmid GNAQ-T2A-GFP construct incorporating all expression cassettes (**Strategy A**), and a multi-plasmid approach designed to avoid C-terminal modification of GNAQ by the T2A cleavage site while improving electroporation efficiency through the use of smaller individual plasmids (**Strategy B; see Methods**).TEAZ applied to adult zebrafish skin (TEAZ-Skin) induced melanoma in wild-type animals using either strategy, with Strategy B significantly reducing tumor latency (**Figure 1C–D, Figure S1A**). Melanomas developed exclusively from clonal expansion of cells harboring edits introduced by all gRNAs. Neither oncogenic GNAQ expression nor individual tumor suppressor loss alone was sufficient to drive tumorigenesis, demonstrating that combined oncogenic signaling and inactivation of all gRNA-targeted tumor suppressors were required for transformation (**Figure S1A**). Consistent with prior reports, loss of *mitfa* markedly accelerated tumor onset following TEAZ-Skin, with tumors arising within ∼30 days in *nacre* (*mitfa^w2/w2^*) and *casper* (*mitfa^w2/w2^*, *mpv17^a9/a9^*) zebrafish (**Figure 1C–D, Figure S1B–D**). In these backgrounds (**see Methods**), oncogenic GNAQ alone was sufficient to drive melanoma, consistent with increased susceptibility of Mitfa-independent cells. Tumor identity was distinguished from xanthomas by gross morphology and pigmentation (**Figure S1E**), and CRISPR-mediated editing of target genes was confirmed by Sanger sequencing (**Figure S2**). Together, these data demonstrate that TEAZ-Skin efficiently induces GNAQ-driven melanoma in adult zebrafish, with Strategy B providing faster tumor onset than Strategy A and *mitfa* loss substantially accelerating disease compared to TEAZ-Skin in wild-type zebrafish (**Figure 1C–D**).

**Figure 1:**
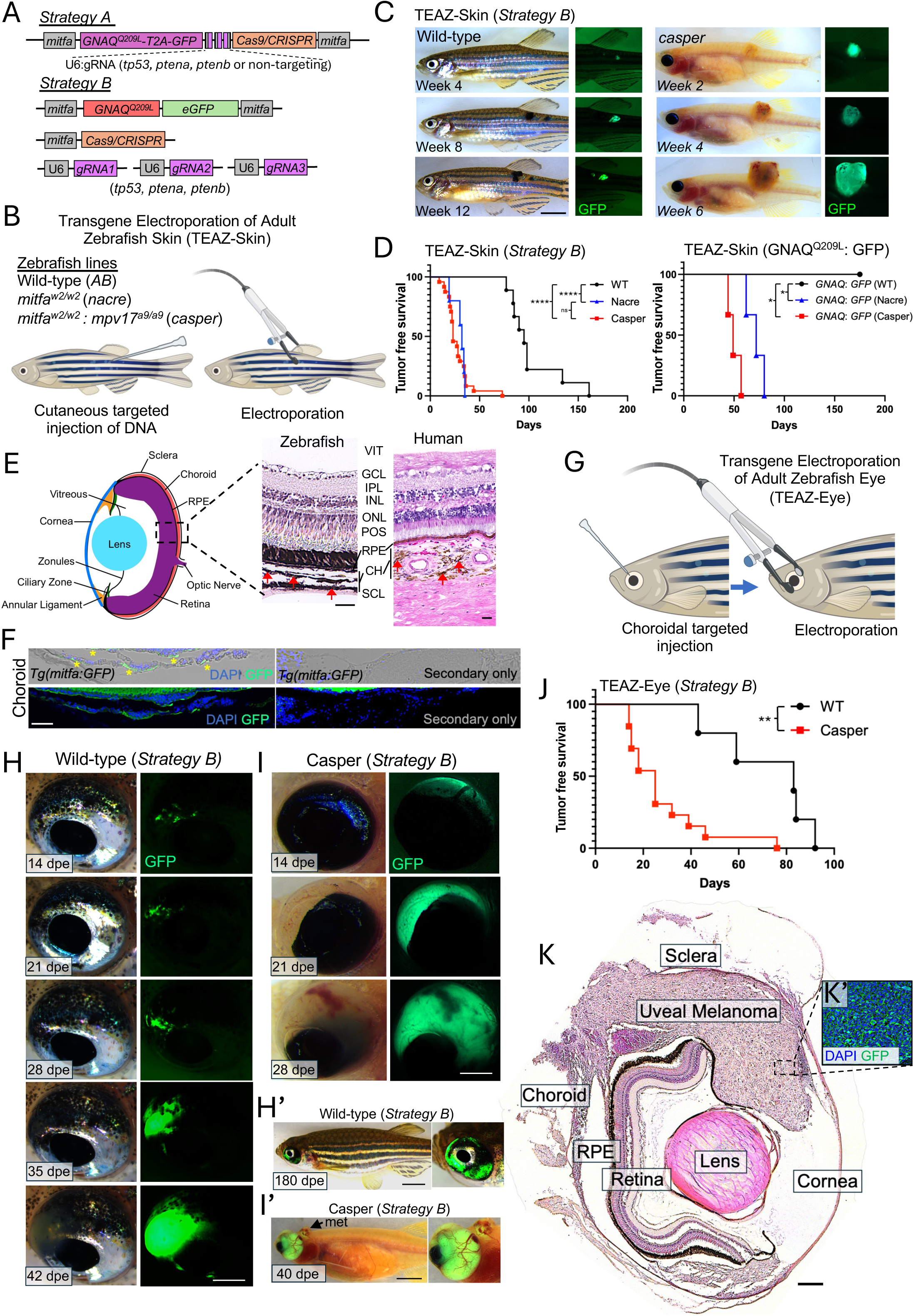
*GNAQ*^Q209L^-induced melanoma by TEAZ-Skin and TEAZ-Eye. (A) TEAZ-Skin was used to induce GNAQ-positive melanoma using two strategies. <u>Strategy A</u> involves a single plasmid construct containing expression cassettes driven by the *mitfa* promoter for GNAQ^Q209L^-T2A-eGFP and Cas9, as well as a U6 promoter driving the expression of three separate gRNAs (targeting *tp53*, *ptena*, and *ptenb*, or a non-targeting control). <u>Strategy B</u> uses a multi-plasmid approach with five plasmids for TEAZ-Skin. Expression cassettes include: (a) *mitfa*:*GNAQ*^Q209L^-*mitfa*:eGFP, (b) *mitfa*:Cas9, (c) *U6*:gRNA (*tp53*), (d) *U6*:gRNA (*ptena*), and (e) *U6*:gRNA (*ptenb*). (B) Schematic of TEAZ-Skin using wild type (*AB*), *nacre* (*mitfa^w2/w2^*) and *casper* (*mitfa^w2/w2^: mpv17^a9/a9^*) zebrafish lines. (C) TEAZ-Skin using Strategy B plasmids in adult wild-type (*AB*) and *casper* zebrafish. Representative brightfield and fluorescent images shown at 4, 8, and 12 weeks post electroporation for WT and at 2, 4, and 6 weeks for *casper*. Scale bar = 0.5 cm. (D) Kaplan-Meier curves were used to estimate tumor-free survival, defined as the time from electroporation until the first visible appearance of GFP+ nodular tumors, in adult zebrafish via TEAZ-Skin. Strategy B wild-type (n=9), *nacre* (n=5), and *casper* (n=24) curves and TEAZ-Skin (*GNAQ*^Q209L^:GFP) curves (n=3 per condition) compared pairwise with logrank (Mantel-Cox) test. ns = not significant, * p-value<0.05; ** p-value<0.01; ***p-value<0.001; ****p-value<0.0001. (E) Schematic of the zebrafish eye with dashed box indicating the retina and choroid. Comparative H&E of the retina and choroid from a wild-type zebrafish (left) and human donor (right) (20×). Choroidal melanocytes indicated with red arrows. VIT, vitreous; ILM, internal limiting membrane, GCL, ganglion cell layer; IPL, inner plexiform layer, INL, inner nuclear layer; ONL, outer nuclear layer; POS, photoreceptor outer segments; RPE, retinal pigmented epithelium; CH, choroid; SCL, Sclera. Scale bars = 20 μm. (F) Immunofluorescent analysis of transgenic *Tg(mitfa:GFP)* zebrafish using an anti-GFP antibody. Sections were counterstained with DAPI to visualize nuclei. Representative image showing GFP expression in *mitfa*-positive cells. Secondary-only antibody was used as background control. Scale bar = 50 μm. (G) Schematic of TEAZ-Eye using choroidal targeted injection. (**H-H’**) Live imaging of adult wild-type zebrafish following TEAZ-Eye with Strategy B plasmids to induce uveal melanoma. Brightfield imaging and the corresponding GFP fluorescence images highlighting tumor progression. Timepoints in day post-electroporation (dpe) as labeled. (H) Scale bar = 250 μm. (H’) Scale bar = 0.5 cm. (**I-I’**) Live imaging of adult *casper* zebrafish over 21 days post-electroporation (dpe) using TEAZ-Eye with Strategy B plasmids to induce uveal melanoma. Brightfield imaging and the corresponding GFP fluorescence images highlighting tumor progression. (I) Scale bar = 250 μm. (I’) Scale bar = 0.5 cm. (**J**) Kaplan-Meier curves comparing tumor-free survival among wild-type (n=5) and *casper* (n=13) zebrafish following TEAZ-Eye induction using Strategy B plasmids. Curves compared pairwise with logrank (Mantel-Cox) test. ** p-value<0.01. (**K, K’**) H&E analysis of uveal melanoma in *casper* zebrafish induced by TEAZ-Eye. Images were acquired at 20× magnification and stitched from 85 fields using Thunder imaging software. Normal eye tissue and uveal melanoma as labeled. Scale bar = 100 μm. (**K’**) Representative 40x magnification image showing GFP expression in tumors cells after anti-GFP immunofluorescent analysis of TEAZ-Eye induced uveal melanoma in *casper* zebrafish.

### The *mitfa* promoter is active in choroidal melanocytes

We next sought to adapt the TEAZ approach to induce melanocyte transformation in the zebrafish eye (TEAZ-Eye). The zebrafish and human eye share highly conserved overall organization and cellular composition (reviewed in ^20^), including distinct retinal layers, retinal pigmented epithelium, and a melanocyte-containing choroidal layer. Most major human ocular compartments have clear zebrafish counterparts, although the zebrafish lacks a fovea and has a proportionally thinner choroid (**Figure 1E**). We first assessed specificity of the *mitfa* promoter in driving transgene expression in choroidal melanocytes by performing immunofluorescent imaging with an anti-GFP antibody in eyes harvested from transgenic *Tg(mitfa:GFP)* zebrafish.

Choroidal melanocytes exhibited strong GFP positivity compared to the secondary antibody-only control (**Figure 1F**). Quenching autofluorescence signal from RPE improved the signal-to-noise ratio (**Figure S3A-B**). We next applied TEAZ-Eye by injecting and electroporating plasmids containing *mitfa* promoter-driven *GFP* expression (*mitfa:GFP*) into the choroid. Eyes were harvested two weeks post-electroporation to assess GFP expression by immunofluorescence. Cells at the injection site exhibited strong GFP signal compared to the secondary antibody-only control (**Figure S3C**), demonstrating that the *mitfa* promoter is sufficient to drive transgene expression in choroidal melanocytes using TEAZ-Eye.

### TEAZ-Eye induces UM in the choroid of wild-type and *casper* zebrafish

A major limitation in the study of UM onset and progression is the lack of anatomically accurate *in vivo* models. To address this gap, we used the TEAZ-Eye technique to introduce oncogenic GNAQ and tumor suppressor gRNAs into the choroidal space of wild-type zebrafish (**Figure 1G-H**). Remarkably, TEAZ-Eye in wild-type zebrafish promoted transformation of cells within the choroid. Live *in vivo* imaging of TEAZ-Eye enabled identification of individual GFP+ cells at the injection site, serving as a proxy for *GNAQ^Q209L^* expression (**Figure 1H-H’**). We next applied TEAZ-Eye to *mitfa*-deficient *casper* zebrafish (**Figure 1I-I’**). As with TEAZ-Skin, tumor latency of TEAZ-Eye injected *casper* zebrafish (n=13) was significantly reduced compared to wild-type (n=5) (**Figure 1J**). H&E analysis confirmed that TEAZ-Eye induced UM within the choroid, progressing into surrounding ocular structures (**Figure 1K**). Notably, 100% (n=10) of tumors formed between the sclera and RPE, highlighting the model’s ability to induce anatomically accurate UM. Finally, anti-GFP immunofluorescent analysis confirmed expression of plasmid constructs following TEAZ-Eye (**Figure 1K’**).

### UM transformation involves activation of neural crest transcriptional programs

During embryonic development, choroidal melanocytes are derived from the cranial neural crest (CNC).^21^ Since our model can detect the earliest events in UM transformation, we asked whether reactivation of CNC programs is one of the initial steps in UM development. To address this point, tumors were derived in wild-type zebrafish using TEAZ-Eye with Strategy B plasmids. At 50-90 dpe, when gross nodular tumors were identified, surgical enucleation was performed, and dissociated cells from the entire eye were subjected to single-cell RNA sequencing (scRNA-Seq). We compared pooled wild-type tumors (2 sequencing replicates) with tumor-free paired eyes or sibling-matched control eyes (2 and 4 pooled eyes, respectively, 2 sequencing replicates) (**Figure 2A, Figure S4**). Given that whole eyes were used as control samples and melanocytes are most abundant in the choroid, melanocytes identified under control conditions are hereafter referred to as choroidal melanocytes. Tumor cells were identified by expression of *GFP* and oncogenic *GNAQ* (**Figure 2B**) whereas primary choroidal melanocytes were marked by expression of *mlana* (**Figure S5**). Annotation of additional cell types within the TME was accomplished by comparing to previously published human choroid and retina scRNA-Seq datasets,^14,22^ the Zebrafish Information Network (ZFIN),^23^ and from the zebrafish embryo single cell atlas “Daniocell”^13^ (**Figure 2A**). Interestingly, tumor cells aligned most closely with choroidal melanocytes, ruling out the retinal pigmented epithelium (RPE) as an origin of our TEAZ-Eye induced tumors (**Figure 2C**). Additionally, profiling *GFP/GNAQ^Q209L^*+ UM cells revealed that tumors exhibited strong expression of neural crest markers (*tfap2a, tfap2c, pax7b, sox10, foxd3*) and melanocytic markers (*mitfa, tfap2e, kita, mlana, pmela, slc22a7a, mlpha, mtbl*) (**Figure 2D**). Unexpectedly, CNC marker genes *twist1* or *dlx1*^24,25^ were not enriched in UM tumors compared to TME. Pathway analysis showed that genes enriched in TEAZ-Eye-derived tumors were strongly associated with MITF-M-dependent gene expression, RAF/MAP kinase, mTOR, PI3K/AKT signaling, NF-kB, RHO GTPase, and NOTCH activity, as well as fatty acid oxidation pathways (**Figure S6A-B**).

**Figure 2:**
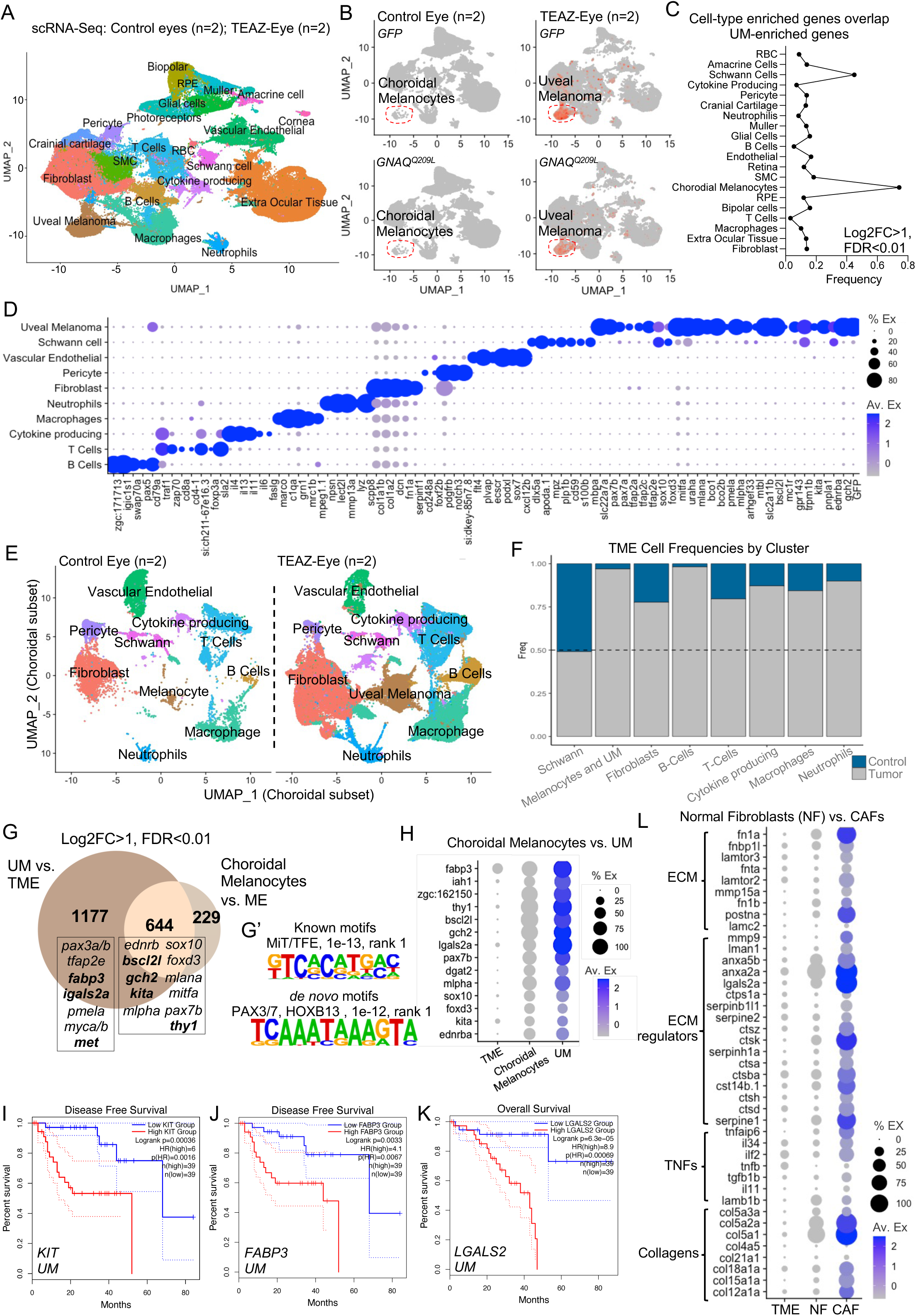
Single cell profiling of uveal melanoma and the TME in adult zebrafish. (A) UMAP representation of single-cell RNA-Seq (scRNA-Seq) from three dissociated wild-type eyes harboring uveal melanoma (UM). UM tumors were generated by TEAZ-Eye (sequencing replicates, n=2). Control eyes represent pools of two, tumor-free paired wild-type eyes and four sibling-matched wild-type eyes (sequencing replicates, n=2). Annotated cell clusters as labeled. (B) UMAP and feature plot showing *eGFP* and *GNAQ*^Q209L^ expression in control eyes and UM tumor. Red dotted line highlights the melanocyte and UM clusters. (C) Comparison of UM tumor marker genes with the markers of cell types within the control eye microenvironment. Only genes with a log2FC > 1 and FDR < 0.01 were included in the analysis. (D) Dot plot representing enriched genes (log2FC > 1; q-value < 0.01) in choroidal cell populations, UM tumors cells, immune cell populations, fibroblasts, Schwann cells, pericytes, and vascular endothelial cells as shown. Dot size indicates the percentage of cells expressing each gene; color intensity (grey to blue) reflects normalized average expression levels (low to high). (E) UMAP clustering of cells in the choroidal microenvironment. Annotated cell clusters as labelled. (F) Comparison of relative cluster sizes between the two conditions, control and TEAZ-Eye, shown in (E). Schwann cells serve as a control cluster, displaying no change in frequency between conditions. Gray = TEAZ-Eye, Blue = Control. (G) Venn diagram representing the overlap of genes between UM tumors and choroidal melanocytes compared to the cells in their respective microenvironments. Enriched genes for each condition were identified by differential expression analysis comparing melanocytes to the microenvironment (ME) or tumor cells to the tumor microenvironment (TME), using a threshold of log2FC > 1 and FDR < 0.01. Genes of interest have been labelled in their respective subsets, bolded genes represent genes associated with worse overall survival or disease-free survival in human UM TCGA datasets. (G’) HOMER promoter motif enrichment analysis of genes enriched in UM tumors vs TME. The top ranked motif for “known” and “*de novo”* motifs are shown with their corresponding p-values. (H) Dot plot showing enriched genes (log2FC > 1; FDR < 0.01) in UM tumor cells, choroidal melanocytes, or the tumor microenvironment (TME). Dot size indicates the percentage of cells expressing each gene; color intensity (grey to blue) reflects normalized average expression levels (low to high). (**I-K**) Kaplan–Meier disease-free survival and overall survival curves generated using uveal melanoma TCGA data. Patients stratified by relative expression levels of *KIT*, *FABP3*, or *LGALS2*. Gene expression thresholds were defined by median expression within each cohort. Log rank p-values as shown. Dotted lines represent 95% confidence interval. (**L**) Dot plot showing enriched genes (log2FC > 1; FDR < 0.01) in cancer-associated fibroblasts (CAF), normal fibroblasts (NF), or the tumor microenvironment (TME) without CAF. Dot size indicates the percentage of cells expressing each gene; color intensity (grey to blue) reflects normalized average expression levels (low to high). Genes have been grouped according to their labelled functions.

Re-clustering of choroidal cells from control and tumor conditions revealed a marked expansion of tumor cells relative to melanocytes, accompanied by increases in immune cells and fibroblasts (**Figure 2E-F**). Immune cell populations within the TME included B-cells (*igic1s1, pax5, cd79a, zgc:153659*), T-cells (*traf1, si:ch211-67e16.3, cd27, zap70, cd8a, sla2*), a population of cytokine producing cells, likely representing T-cells (*il4, il13, il11, ca2, il34, il6*) as well as macrophages (*marco, ccl34a.4, mrc1b, c1qa/b/c, grn1*) and neutrophils (*cpa5, lect2l, mmp9, npsn, scpp8*). Additional cell types within the choroidal TME included fibroblasts (*fn1a*, *col1a1a, col1a1b, col1a2, col5a1, dcn, serpinf1*), pericytes (*pdgfrb, notch3, foxf2b, cd248a*), Schwann cells (*dlx5a, apoda.1, mpz, plp1b, cd59, s100b, mbap*) and vasculature endothelial cells (*vwf, flt4, epas1b, ecscr, fabp11a*) (**Figure 2D-F**, **Table S1**).

We next compared choroidal melanocytes and UM tumors, revealing a significant overlap in gene expression (n=644; hypergeometric p<0.0001) alongside a large set of tumor-enriched genes (n=1177) (**Figure 2G**). Shared genes included melanocytic regulators (*ednrb, sox10, foxd3, mitfa, mlpha, kita, mlana, pax7b*), supporting a melanocytic origin, with many transcripts further upregulated in UM tumors compared to choroidal melanocytes (**Figure 2H**). HOMER promoter motif enrichment analysis of genes enriched in UM tumors revealed binding motifs for the MiT/TFE family of transcription factors, as well as PAX3/7 and HOXB13, suggesting that these regulators may contribute to UM-specific transcriptional programs (**Figure 2G’**).

To assess the relevance of our model to human disease, we examined whether genes enriched in zebrafish UM tumors were associated with clinical outcomes in patients. Differential expression analysis identified a gene signature, including *LGALS2, FABP3/5, GCH, MET*, and *KIT*, whose high expression was associated with reduced overall and disease-free survival and with TCGA molecular subtypes 3 and 4,^26^ corresponding to aggressive UM (**Figure 2G, 2I-K, Figure S7-8, Table S2**).

### Cancer associated fibroblasts within the uveal melanoma TME upregulate components of the extracellular matrix (ECM)

Recent studies have significantly advanced our understanding of the TME as an active driver of melanoma progression.^27,28^ While it is well established that components of the TME, particularly cancer-associated fibroblasts (CAFs), play a key role in malignancy,^29–31^ *in vivo* models to study such interactions in UM remain limited. Our model allows for direct comparison of the microenvironment in TEAZ-Eye and control injected eyes within the same animals (i.e., left and right eye). We compared the transcriptome of CAFs and normal fibroblasts in wild-type zebrafish. Upon re-clustering and differential gene expression analysis, we found that several ECM proteins were upregulated in CAFs, indicating ECM remodeling in the presence of UM. Enriched genes broadly fell into four categories: ECM, ECM regulators, TNFs, and collagens.

Notably, fibronectin (*fn1a*) was among the most significantly upregulated genes in CAFs compared to normal fibroblasts (q<0.0001). Several collagens were also found to have higher expression in CAFs, including Collagen I (*col1a1b, col1a2),* Collagen V (*col5a1, col5a2a, col5a3a)* and Collagen XII (*col12a1a)* (**Figure 2L, Table S3**). Consistently, cell–cell communication analysis using CellChat^32^ identified FN1 and collagen signaling from CAFs to tumor cells as significantly enriched pathways and predicted the transmembrane heparan sulfate proteoglycan SDC4 as a mediator of these interactions in UM cells (**Figure S9**).

To control for injection-related fibrosis potentially causing fibroblast activation and upregulation of ECM components, we mock-injected three wild-type eyes with plasmids containing *mitfa*:GFP and *mitfa*:Cas9 cassettes followed by electroporation (mock-TEAZ) and performed scRNA-Seq. Fibroblasts from mock-injected sibling controls (three pooled eyes), non-injected sibling controls (four pooled eyes), and non-injected tumor-matched control eyes (two pooled eyes) were equally distributed on the UMAP, with no alteration in *fn1a* or *lgals2a* expression or other changes to extracellular matrix (ECM) genes upon mock-TEAZ (**Figure S10**). These findings indicate that TEAZ-Eye tumor induction depends on genetic modification of target cells, not simply physical injection, triggering a CAF response that may be critical to melanoma progression.

### Skin and eye primary melanocytes, and their resulting tumors, have unique gene expression profiles

Having established a system to induce genetically identical tumors in the eye and skin of adult zebrafish, we asked whether tumor transcriptional programs differed by anatomical site. Such differences could reflect site-specific tumor–stromal interactions or intrinsic differences between uveal and cutaneous melanocytes. We generated TEAZ-Eye and TEAZ-Skin tumors using Strategy B plasmids and performed scRNA-Seq with matched control eye and skin tissues (**Figure 3A**). We compared tumors to a combined TME and performed a parallel analysis of eye and skin primary melanocytes relative to the combined microenvironment (ME) (**Figure 3B, Table S4**). *GNAQ^Q209L^* expression levels were comparable between eye and skin wild-type tumors (**Figure S11A**) and DEGs showed substantial overlap between sites, with 41% of 2,192 genes shared, consistent with common transcriptional programs driven by oncogenic GNAQ during transformation (**Figure 3B**). Notably, only 25.8% of DEGs overlapped between eye and skin melanocytes, indicating substantial transcriptional differences in melanocytes originating from distinct anatomical regions (**Figure 3B**).

**Figure 3:**
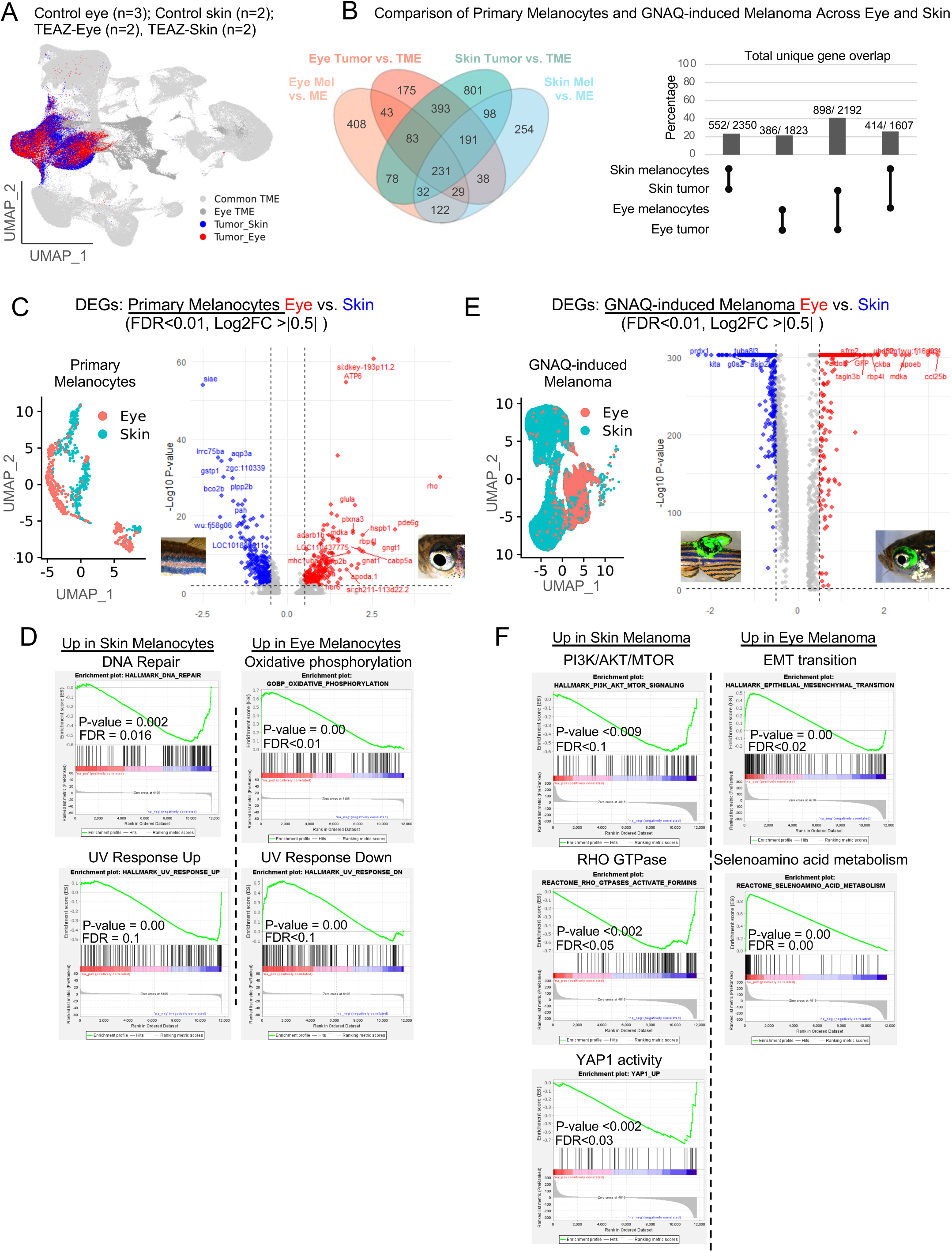
Comparison of GNAQ-driven tumors and primary melanocytes in the skin and eyes of adult zebrafish. (A) UMAP representation of GNAQ-positive tumors generated using TEAZ-Eye and TEAZ-Skin, along with their respective control tissues using scRNA-Seq. Tumor samples include three dissociated wild-type eyes harboring UM tumors generated via TEAZ-Eye and two dissociated wild-type skin UM tumors generated via TEAZ-Skin (n=2 sequencing replicates each). Control eye samples consist of a pool of two uninjected tumor paired wild-type eyes, three mock-injected sibling-matched wild-type eyes, and four uninjected sibling-matched wild-type eyes (n=3 sequencing replicates). Control skin samples consist of dissociated normal skin from two wild-type zebrafish (n=2 sequencing replicates). Cell clusters are annotated as tumor cells (red = Eye; Blue = Skin), tumor microenvironment (TME) cells shared by both eye and skin samples (light gray), and TME cells specific to the eye samples (dark gray). (B) Venn diagram showing overlapping genes between TEAZ-Eye, TEAZ-Skin, choroidal melanocytes, and skin melanocytes compared to the cells in their respective microenvironments. Enriched genes for each condition were identified by differential expression analysis comparing melanocytes to the microenvironment (ME) or tumor cells to the tumor microenvironment (TME), using a threshold of log2FC > 0.5 and FDR < 0.01. Bar chart displaying overlaps between enriched gene sets from (B). Percentages represented as the number of overlapping genes over the total unique genes in both gene sets. (C) UMAP representation of primary melanocytes from both eye (salmon) and skin (teal). Volcano plot of differentially expressed genes (DEGs) between eye and skin primary melanocytes (log2FC > |0.5| and FDR < 0.01). (D) GSEA enrichment plots of pathways that were enriched in skin melanocytes and of pathways enriched in eye melanocytes from (C). P-values and FDRs as labelled. (E) UMAP representation of GNAQ^Q209L^-induced melanoma from both eye (salmon) and skin (teal). Volcano plot of differentially expressed genes (DEGs) between eye and skin tumors (log2FC > |0.5| and FDR < 0.01). (F) GSEA enrichment plots of pathways that were enriched in skin melanoma and of pathways enriched in eye melanoma from (E) P-values and FDRs as labelled.

We hypothesized that DEGs shared by only the two tumor sites and not the primary melanocytes would represent essential mechanisms for UM growth and survival. Consistent with this hypothesis, pathway analysis revealed enrichment of pathways involved in cancer proliferation and metabolism (**Figure S12A**). Additional enriched pathways included WNT signaling, PTEN regulation, and RAF/MAPK signaling, which have been previously implicated in UM pathogenesis.^33–41^ In contrast, IPA analysis of gene sets shared between eye and skin melanocytes represented homeostatic pathways including enrichment of MITF-M–dependent programs, as well as pathways related to nucleotide biosynthesis and mTORC1-regulated amino acid metabolism (**Figure S12B**).

We next re-clustered normal melanocytes from eye and skin for direct comparison, revealing distinct UMAP clusters (**Figure 3C**). We identified 186 DEGs enriched in eye melanocytes and 229 DEGs enriched in skin melanocytes (log2FC > |0.5|, FDR< 0.01), while 3,387 genes were not differentially expressed (FDR > 0.01, expressed in ≥10% of cells, **Table S5**). GSEA revealed that skin melanocytes were enriched for DNA repair pathways and UV response-associated genes (**Figure 3D**). In contrast, eye melanocytes showed enrichment for pathways involved in UV response downregulation and oxidative phosphorylation. We next re-clustered and directly compared eye and skin tumors, which also separated by tissue of origin on the UMAP (**Figure 3E**). We identified 230 DEGs enriched in eye tumors and 375 DEGs enriched in skin tumors (log2FC > 0.5, FDR < 0.01), while 5,754 genes were not differentially expressed between tumor types. Eye tumors were enriched for epithelial-to-mesenchymal transition (EMT) and selenoamino acid metabolism pathways (**Figure 3F**). In contrast, skin tumors were enriched for PI3K–AKT–mTOR signaling and RHO GTPase–mediated activation and YAP1 activity. Together, these analyses demonstrate that both primary melanocytes and GNAQ^Q209L^-driven melanomas exhibit site-specific transcriptional programs.

### Melanocyte progenitor cells are highly susceptible to oncogenic GNAQ transformation in *mitfa*-deficient zebrafish

H&E analysis of *nacre* and *casper* zebrafish eyes showed loss of choroidal melanocytes (**Figure 4A**), whereas melanin in the RPE remained less affected, as previously demonstrated.^42^

**Figure 4:**
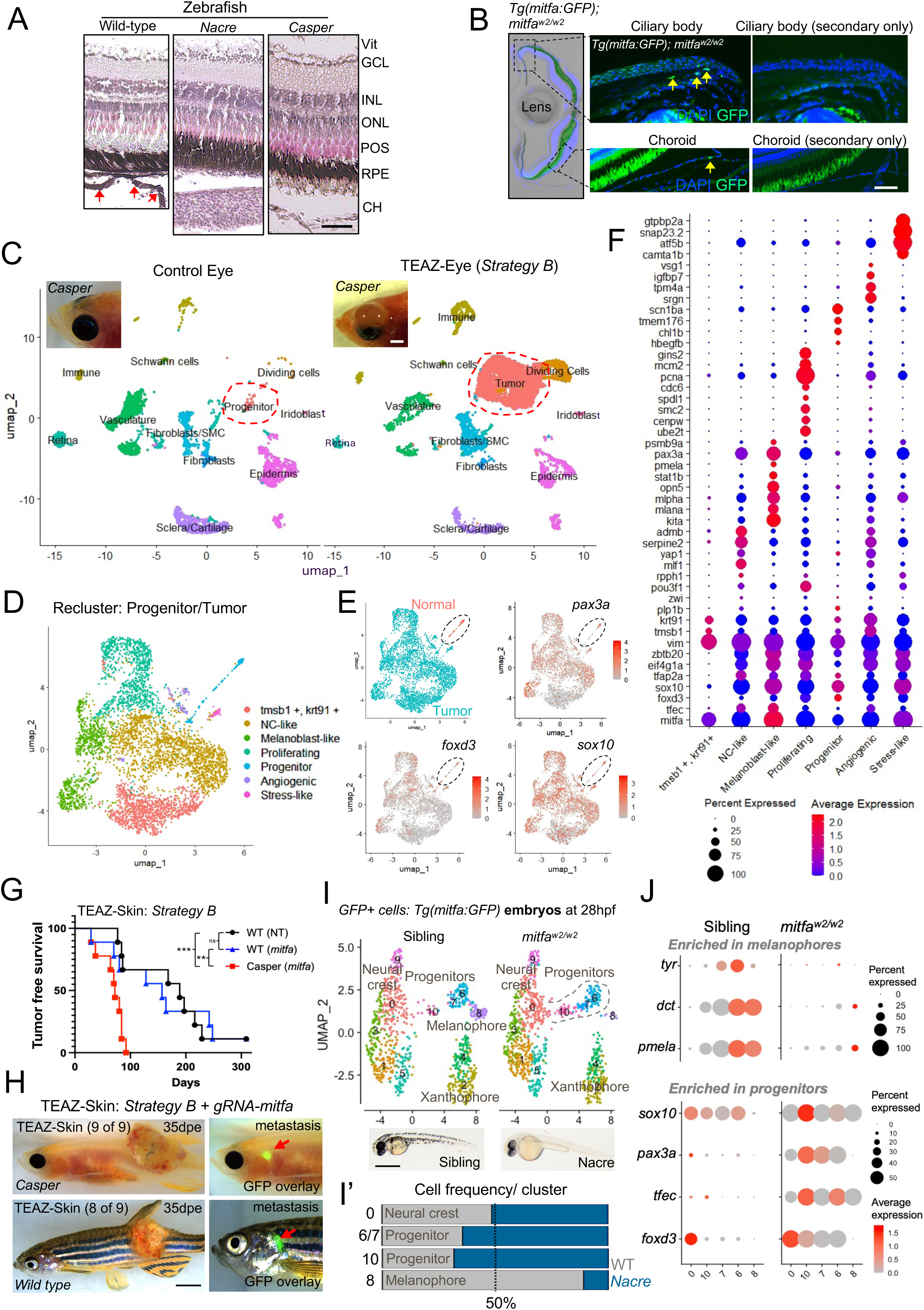
*mitf*-independent melanocyte progenitor cells are expanded in *mitfa*-deficient zebrafish and are susceptible to *GNAQ*^Q209L^ transformation. (A) H&E analysis representation of the retina and choroid in wild-type (left), *nacre* (middle), and *casper* (right) zebrafish (20x magnification). Note the loss of melanocytes (red arrows) in the choroid of *nacre* and *casper* eyes. “Vit” = vitreous, “GCL” = ganglion cell layer, “INL” = inner nuclear layer, “ONL” = outer nuclear layer, “POS” = photoreceptor outer segment, “RPE” = retinal pigmented epithelium, “CH” = choroid. Scale bar = 50 μm. (B) Immunofluorescent analysis of transgenic *Tg(mitfa:GFP); mitfa^w2/w2^* (*nacre*) zebrafish using an anti-GFP antibody. Sections were counterstained with DAPI to visualize nuclei. Representative images showing GFP-positive cells in the ciliary body and choroid are shown by yellow arrow heads. Secondary-only antibody was used as background control. Scale bar = 20 μm. (C) UMAP representation of single-cell RNA sequencing (scRNA-Seq) data from three paired dissociated *casper* eyes and from three *casper* eyes harboring uveal melanoma (UM) tumors induced using Strategy B via TEAZ-Eye (n=1, sequencing replicate). Annotated cell clusters are labeled. Progenitor cells and UM tumor clusters are outlined with dashed red lines. Representative brightfield images of a normal eye and a UM tumor in *casper* zebrafish are shown in the top left corner of the UMAP. Scale bar = 250 μm. (D) Re-clustered UMAP representing progenitor cells from control eyes and uveal melanoma (UM) tumor cells from TEAZ-Eye–injected *casper* zebrafish. Heterogeneous cell populations as labeled. (E) Re-clustered UMAP from (D) showing melanocyte progenitor cells from control eyes in red and UM tumor cells from TEAZ-Eye in blue. Feature plots for *pax3a, foxd3* and *sox10* expression are shown, dashed line marks the progenitor cell population. (F) Dot plot illustrating differentiational expressed genes (log2FC > 0.5; FDR < 0.05) between heterogeneous cell populations in (D). Dot size indicates the percentage of cells expressing each gene; color intensity (blue to red) reflects normalized average expression levels (low to high). (G) Kaplan–Meier curves comparing tumor-free survival in wild-type and *casper* zebrafish following TEAZ-Eye injection using Strategy B plasmids for *mitfa* conditional knockout. Wild-type zebrafish were injected with either Strategy B plasmids plus non-targeting gRNAs (n=9) or Strategy B plasmids plus *mitfa*-targeting gRNAs (n=9). *Casper* zebrafish were injected with Strategy B plasmids and *mitfa*-targeting gRNAs (n=9). (H) Representative brightfield and GFP-overlay images of tumors that formed in *casper* (9 of 9) and wild-type zebrafish injected with *mitfa*-targeting gRNAs (8 of 9). Metastatic cells are indicated by red arrows. Scale bar = 0.5 cm. (**I, I’**) UMAP obtained after clustering GFP-positive cells sorted from *Tg(mitfa:GFP)* and *Tg(mitfa:GFP); mitfa^w2/w2^* (*nacre*) zebrafish embryos at 28 hours post fertilization (hpf). Annotated cell clusters as labelled. (**I’**) Comparison of relative cluster size for the two genotypes; *Tg(mitfa:GFP)* and *Tg(mitfa:GFP); mitfa^w2/w2^* (*nacre*) in (I). Scale bar = 0.5 mm. (**J**) Dot plot illustrating DEGs (log2FC > 0.5; FDR < 0.05) in melanophores and progenitor cells in *mitfa*^w2/w2^ and sibling embryos at 28hpf. Dot size indicates the percentage of cells expressing each gene; color intensity (grey to red) reflects normalized average expression levels (low to high).

These observations imply that an *mitfa*-independent melanocyte progenitor cell persists in *nacre* and *casper* eyes and that they are susceptible to oncogenic GNAQ-driven transformation. To identify such cells, we crossed *nacre* (*mitfa^w2/w2^*) with transgenic *Tg(mitfa:GFP)* lines to create *Tg(mitfa:GFP); mitfa^w2/w2^*zebrafish. We then harvested adult eyes at 5 months and performed anti-GFP immunofluorescence. While GFP+ melanocytes were readily detected in the choroid of *Tg(mitfa:GFP)* zebrafish (**Figure 1F, Figure S3**), staining of *Tg(mitfa:GFP); mitfa^w2/w2^* eyes revealed rare cells with GFP-positivity (**Figure 4B**). Such GFP+ cells were primarily located in the ciliary body, with isolated cells also found in the choroid, suggesting the persistence of an *mitfa*-independent progenitor population in adult *mitfa*-deficient zebrafish eyes (**Figure 4B**). We next asked whether such cells retain the ability to undergo melanocyte differentiation by using TEAZ-Eye to deliver plasmids containing the *mitfa*-promoter driving expression of *mitfa* (i.e., to overexpress *mitfa* mRNA). Interestingly, re-expression of *mitfa* transcript resulted in expansion of choroidal melanocytes in wild-type animals and rescue of choroidal melanocytes in *mitfa*-deficient (*nacre*) zebrafish as shown by H&E analysis (**Figure S13A-B’**). To assess whether GFP-positive cells are sensitive to oncogenic transformation by GNAQ, we expressed *mitfa*:*GNAQ^Q209L^* in *Tg(mitfa:GFP); mitfa^w2/w2^* zebrafish using TEAZ-Eye. In all injected fish (n=3), UM tumors were GFP-positive, suggesting they originated from GFP+ progenitor cells within the eye of *Tg(mitfa:GFP); mitfa^w2/w2^* zebrafish (**Figure S13C**).

### Melanocyte progenitor cells express neural crest marker genes in *mitfa*-deficient zebrafish

We performed scRNA-Seq and pseudotime analysis on 3 pooled UM tumors from the eyes of *casper* zebrafish and compared them to the pooled tissue from 3 normal *casper* eyes (**Figure 4C, Figure S14**). Within the *casper* eyes, we identified a population of cells expressing the *mitfa^w2/w2^* allele that clustered closely with UM tumors under TEAZ-Eye conditions, termed progenitors (**Figure S15A-B and Tables S6-8**). We re-clustered the progenitor and tumor cells (**Figure 4D**), revealing seven distinct clusters consisting of neural crest-like (NC-like), melanoblast-like, proliferating, progenitor, angiogenic, stress-like, as well as an unidentified cluster marked by *tmsb1, krt91* and *vim* (**Figure 4D, Table S9**). Unsupervised pseudotime analysis supports a lineage trajectory from the progenitor cells to heterogeneous UM tumor cells (**Figure S15C**). Notably, the progenitor cell population in normal *casper* eyes expressed neural crest and melanocyte stem cell marker genes, including *tfap2a, foxd3, sox10*, *pax3a, vim,* and *yap1* (**Figure 4E-F**). UM tumor cells retained expression of these genes but activated additional genes associated with pigment progenitor cell function, including *tfec* and *kita* (**Figure 4F**).

Notably, *mitfa* expression in tumors and progenitor cells in *casper* zebrafish reflects the expression of the mutant *mitfa^w2/w2^*allele, which functions as a lineage tracer to mark cells that would normally express *mitfa* under wild-type conditions (**Figure S15B**). Interestingly, despite the lack of Mitfa activity in *casper* zebrafish, GNAQ^Q209L^-positive UM tumors included a melanoblast-like cluster that exhibited expression of pigmentation genes, including *pmela, mlpha*, and *mlana,* likely explaining the occasional emergence of melanin within tumors as they progress in *casper* zebrafish.

### Conditional deletion of *mitfa* in adult zebrafish fails to recapitulate germline loss-of-function effects on tumor-free survival in GNAQ**^Q209L^**-driven melanoma

We next asked whether the accelerated growth of GNAQ^Q209L^-driven tumors in *mitfa*-deficient zebrafish was attributable to increased oncogene expression or cell-intrinsic loss of Mitfa in adult melanocytes. scRNA-Seq analysis showed higher GNAQ^Q209L^ expression in wild-type tumors than in *casper* tumors (**Figure S11B**), thus excluding oncogene overexpression as the basis for the accelerated phenotype. To directly test cell-intrinsic effects of *mitfa* loss in adult melanocytes, we performed conditional somatic deletion of *mitfa* using Strategy B plasmids with either non-targeting gRNAs or gRNAs targeting exon 5 of *mitfa*. TEAZ-Skin was conducted in three groups: (1) *casper* zebrafish with *mitfa*-targeting gRNAs (n=9; which controls for off-target effects of the gRNA and for potential function of the *mitfa^w2/w2^* allele); (2) wild-type zebrafish with non-targeting gRNAs (n=9); and (3) wild-type zebrafish with *mitfa*-targeting gRNAs (n=9). GFP expression was confirmed in all injected animals (**Table S11)**. As expected, by 100 dpe tumors developed in 100% (9 of 9) of *casper* zebrafish. Interestingly, there was no significant difference in tumor latency between wild-type and *mitfa* conditional-KO, with both indicating significantly longer latency than in *casper* animals (**Figure 4G–H**). Thus, somatic loss of *mitfa* in adult melanocytes is insufficient to reproduce the reduced tumor-free survival observed with germline *mitfa* deficiency. Notably, *mitfa*-deficient tumors arising in either background developed distal metastases, a feature not observed in *mitfa* wild-type TEAZ-Skin tumors (**Figure 4H, Figure S16**). Together, these findings argue against Mitfa as a repressor of GNAQ-induced transformation in adult melanocytes and indicate that somatic loss in the adult is insufficient to drive tumor initiation.

### *mitfa*-deficient zebrafish exhibit expanded *pax3a* and *tfec* positive melanocyte progenitors during embryogenesis

We next asked whether melanocyte progenitor cells are expanded in *mitfa* mutant zebrafish relative to wild-type zebrafish. To test this prediction, we performed scRNA-Seq on GFP-positive cells sorted from *Tg(mitfa:GFP);mitfa^w2/w2^* and *mitfa^+/w2^* sibling-matched transgenic zebrafish embryos at 28 hours post fertilization (hpf), when melanocytes (zebrafish melanophores) begin to differentiate (**Figure 4I, Figure S14**). We assigned cell-type annotations based on our previously annotated GFP-positive cells from *Tg(mitfa:GFP)* embryos.^43^ The 10 clusters included six main cell types: neural crest cells (*sox10, foxd3*), a tripotent precursor of melanoblasts (M), iridoblasts (I), and xanthoblasts (X) (*tfap2a*, *cdkn1ca, slc15a2, ino80e, id3, mycn, tfec*), termed MIX cells, a cluster that expressed high levels of melanoblast/xanthoblast markers (*mitfa, erbb3b, impdh1b, gch2, id3*), termed MX cells, a melanoblast cluster (*mitfa, dct, pmel, tyr*), as well as two additional clusters corresponding to xanthoblasts and xanthophores (**Table S10**). For this analysis we referred to MIX and MX cells as melanophore progenitors (**Figure 4I**). As expected, melanophore cells were reduced in *Tg(mitfa:GFP): mitfa^w2/w2^* zebrafish (**Figure 4I’**). However, melanophore progenitors were expanded in *Tg(mitfa:GFP); mitfa^w2/w2^*, while neural crest cells remained unaffected (**Figure 4I-I’**). Melanophore markers were downregulated in GFP-positive cells sorted from *Tg(mitfa:GFP); mitfa^w2/w2^,* whereas melanocyte progenitor markers (*pax3a, sox10, foxd3*), as we identified in adult zebrafish eyes, were upregulated in embryonic progenitors from *Tg(mitfa:GFP); mitfa^w2/w2^* embryos (**Figure 4J, Table S12**). A role for *pax3a* in adult fish eye tissue has not been previously characterized; however, recent studies have identified *PAX3* expression in ocular surface melanocytes,^44^ and another study reports an association between *PAX3* and stem cell markers in UM progression.^45^

### Transcriptional programs enriched in oncogenic BRAF- versus GNAQ-driven melanoma overlap with specific melanocyte progenitor populations

Oncogenic BRAF and NRAS are unable to promote transformation in the absence of *mitfa,*^6,46^ suggesting that distinct transcriptional programs, potentially reflecting unique cellular origins, are required for UM and CM transformation. To address this hypothesis, we compared previously published scRNA-Seq datasets in transgenic *Tg(mitfa:BRAF^V600E^): tp53^-/-^* : *mitfa^w2/w2^* zebrafish where TEAZ-Skin was used to rescue *mitfa* and induce melanoma,^46^ to our GNAQ^Q209L^-induced TEAZ-Skin tumors (**Figure 5A)**. We predicted that by analyzing primary tumors, the transcriptional signature of the origin cell will be retained and can be identified by overlapping gene signatures identified in the embryonic melanocyte lineage.^43^ Interestingly, while the expression of *mitfa* was significantly higher in GNAQ-positive tumors, the expression of pigmentation genes; including *tyrp1b, dct, scl45a2, sox10* and *pmela* were significantly higher in BRAF-positive melanoma (**Figure S17**). Genes enriched in GNAQ-UM and BRAF-CM were defined as those with a Log2FC > 1 and Adj p-value < 0.0001 between the two tumor types (**Figure 5B, Table S13**). We next overlapped this differential gene signature with genes expressed by cells within the melanocyte lineage isolated from zebrafish embryos (**Figure 5C**). As expected, BRAF-driven tumors strongly mapped to melanoblast and melanophore clusters (**Figure 5D**), supporting recent findings that BRAF-driven melanoma originates from melanoblasts, a melanocyte progenitor population^46,47^. When comparing GNAQ-enriched genes to the melanocyte lineage, we found that signatures were not confined to a single cluster but were distributed among neural crest cells, melanocyte progenitors, and xanthophore clusters (**Figure 5D; Figure S18**). Such results support the hypothesis that GNAQ- and BRAF-driven melanomas depend on distinct transcriptional programs, reflecting less and more differentiated melanocytic states, respectively.

**Figure 5:**
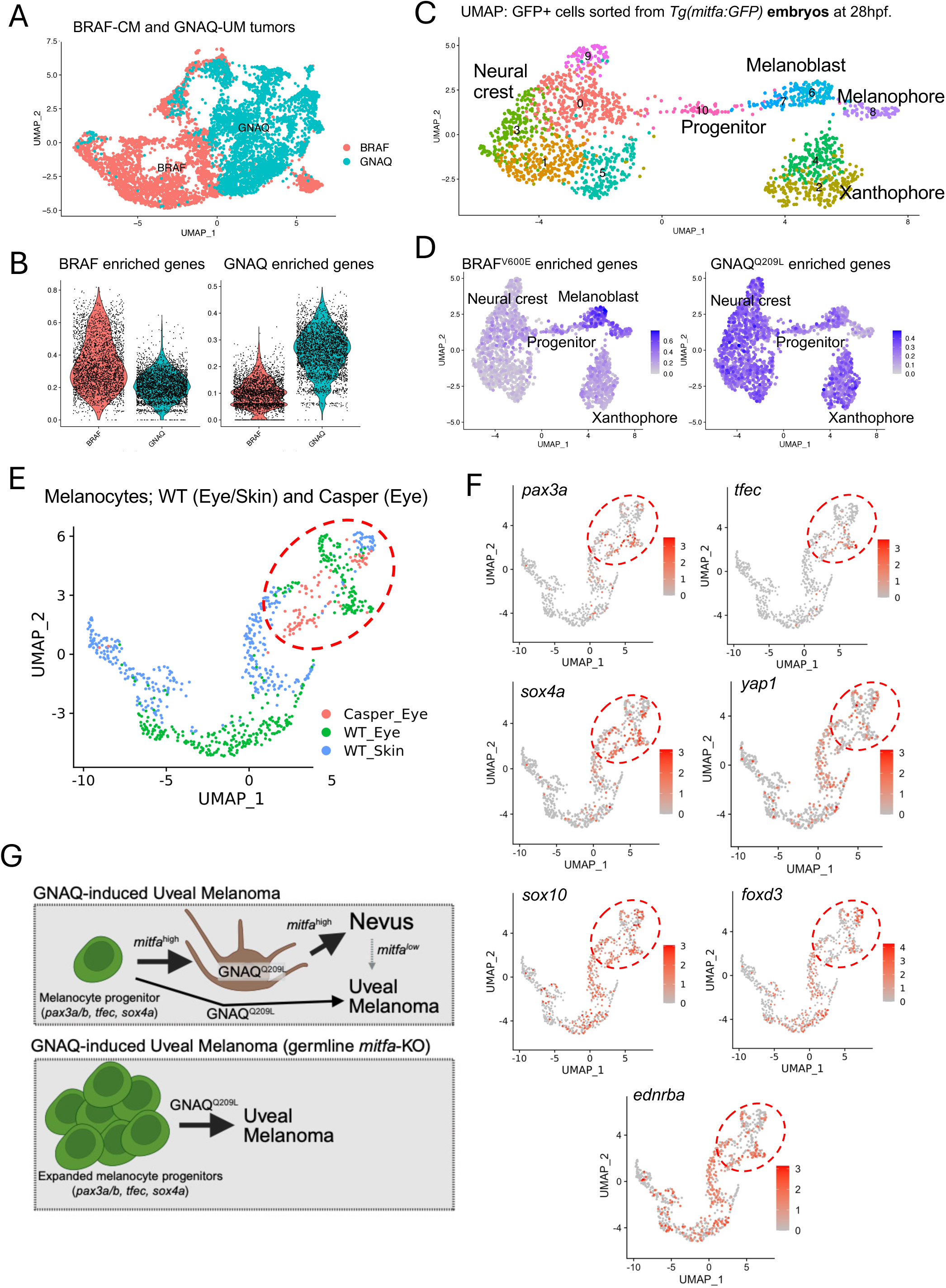
Oncogenic GNAQ and BRAF transform transcriptionally distinct cells within the melanocyte lineage. (A) UMAP obtained after clustering tumor cells from *Tg(mitfa:BRAF^V600E^): tp53^-/-^* : *mitfa^w2/w2^*zebrafish where TEAZ-Skin was used to rescue *mitfa* expression and knockout *ptena* and *ptenb* to induce cutaneous melanoma,^46^ along with our GNAQ-induced tumors by TEAZ-Skin (Strategy B: *mitfa:Cas9, U6:*gRNA*-tp53, U6:*gRNA*-ptena, U6:*gRNA*-ptenb, mitfa:GNAQ*^Q209L^*-mitfa:GFP*). BRAF-induced tumors in red and GNAQ-induced tumors in blue. (B) Violin plot showing differentially expressed genes (FDR < 0.05) between BRAF-induced and GNAQ-induced tumors from (A). The plot displays the distribution of gene expression across individual cells, with the width indicating cell density at specific expression levels and the height representing the range of expression values. (C) Integrated UMAP after clustering GFP-positive cells sorted from *Tg(mitfa:GFP)* and *Tg(mitfa:GFP); mitfa^w2/w2^*(*nacre*) zebrafish embryos at 28 hours post fertilization (hpf). Annotated cell clusters as labelled. (D) Average enrichment of signature genes from BRAF- and GNAQ-driven tumors projected onto the UMAP in (C) using UCell with ’FeaturePlot’ function. Signature genes represent the top 50 enriched genes from each tumor genotype that are also expressed in the embryonic melanocyte lineage. (E) UMAP of primary melanocytes and melanocyte progenitors from 9 control wild-type eyes, 2 control wild-type skin sections, and 3 control *casper* eyes. Cells are labeled according to their genotype (*casper* and wild-type) and tissue origin (eye and skin). (F) Feature plots representing the relative mRNA expression of *pax3a*, *tfec, sox4a, yap1, sox10*, *foxd3* and *ednrba* in melanocytes described in (E). Color intensity (grey to red) reflects normalized average expression levels (low to high). The red dotted line indicates cells that are clustering closely with the melanocyte progenitors identified in *casper* eyes. (G) Schematic illustrating GNAQ^Q209L^-driven uveal melanoma (UM) in the context of tumor suppressor loss. Melanocyte progenitor cells (*pax3a/b, tfec, sox4a*) can directly give rise to UM when oncogenic GNAQ^Q209L^ is present (black arrow), or differentiate into MITF^high^ melanocytes that preferentially form nevi (black arrow). These nevi may subsequently progress to UM following reduced MITF activity (gray dashed arrow). In contrast, germline *mitfa* loss blocks melanocyte differentiation and expands melanocyte progenitors, enabling GNAQ^Q209L^ to drive UM directly from this progenitor population.

We next asked whether less differentiated melanocytes, similar to the *mitfa*-deficient progenitors in *casper* eyes, are also present in adult wild-type eyes. Since melanocyte stem cells are known to reside in the zebrafish skin,^48^ we included skin tissue as a control and re-clustered melanocytes from wild-type skin and eyes with progenitor cells from *casper* eyes. Interestingly, a subset of wild-type melanocytes from both eye and skin samples clustered closely with *casper* progenitors (**Figure 5E, Figure S19**) and expressed pigment cell progenitor marker genes, including *pax3a,*^49^ *tfec,*^42^ *sox10,*^50^ *sox4a,*^51,52^ *foxd3,*^53^ and *yap1* (**Figure 5F**). These findings suggest either melanocyte plasticity within the eye or the presence of an eye-resident melanocyte progenitor cell population in adult wild-type zebrafish. In support of this notion, reanalysis of published scRNA-Seq datasets from human^22^ and mouse^54^ eyes identified *PAX3*-positive melanocyte subpopulations (**Figure S20**) Moreover, we confirmed the presence of PAX3-positive cell populations in adult C57BL/6 mouse eyes by immunohistochemistry (**Figure S21**). Together, these data support the existence of a *PAX3*-expressing melanocyte precursor population in both fish and mammalian eyes, which may mirror *PAX3* expression in skin melanocyte stem cells.^49,55^ Such cells may be more susceptible to GNAQ^Q209L^-driven transformation and represent candidate cells of origin for UM in this context (**Figure 5G**).

## Discussion

Our UM model enables analysis of the earliest events in UM initiation and progression. Compared with transgenic approaches, this system produces anatomically correct tumors with shorter latency. Importantly, genes identified in this model are associated with survival differences in human uveal melanoma, offering an efficient and clinically relevant platform for studying UM pathogenesis.

We found that fatty acid binding proteins, particularly *fabp3*, are enriched in UM tumors. While lipid droplets have been identified as vulnerable targets in CM and regulators of melanoma plasticity,^46,56^ lipid signaling in UM has received limited attention.^57–60^ Interestingly, high expression of *FABP3* and *FABP5* are associated with worse disease-free survival in UM but not CM patients. Our analysis also identified *LGALS2* and *GCH*, two fatty acid metabolism-related genes whose expression was strongly associated with worse overall survival in UM but showed an inverse relationship in CM. We additionally found increased *kita* expression in GNAQ-driven tumors compared to choroidal melanocytes, which is particularly notable given recent studies linking KIT-positive UM tumors with poor prognosis and chromosome 3 monosomy.^61^ In addition, *met* expression was elevated in UM tumors relative to choroidal melanocytes. MET signaling is a key pathway in UM pathogenesis, with established roles in tumor invasion, metastatic progression, and responsiveness to hepatocyte growth factor (HGF), particularly in the context of liver metastasis.^62,63^ Notably, MET activation has also been described in other cancer types through ligand independent fibronectin-integrin α5β1-mediated signaling.^64^ Given the marked increase of *fn1a* in choroidal CAFs, it is plausible that fibroblast-driven ECM remodeling enhances MET signaling in UM. Furthermore, expression of these genes was highest in molecular subclasses 3 and 4 of the UM TCGA dataset which represents the subset of patients with the highest rates of BAP1 loss and metastasis.^26^ Such observations demonstrate the utility of our model in uncovering key genes with disease-specific roles in UM whose function could be further explored through genetic perturbations in zebrafish.

To explore how anatomical context influences transcriptional states, we compared tumors induced by identical genetic perturbations in the skin and eye. We find that transcriptional profiles of UM tumors arising from melanocytes in the choroid and skin are influenced by more than the intrinsic differences of their cells of origin. For example, eye tumors were enriched for EMT programs compared with skin tumors. In contrast, eye melanocytes did not exhibit these signatures relative to skin melanocytes, suggesting that signals from the ocular microenvironment may promote a more mesenchymal state. Alternatively, these differences may represent a unique cellular origin of UM in the eye and skin. Further investigation is needed to determine whether these differences are driven by the microenvironment or by intrinsic properties of melanocytes or precursors cells from distinct anatomical sites.

An interesting distinction between GNAQ and BRAF oncogenes is their differential dependency on Mitfa activity for tumorigenesis. While BRAF^V600E^ requires Mitfa activity to induce melanoma,^6^ GNAQ^Q209L^-driven tumor latency was significantly reduced in *mitfa*-deficient zebrafish compared to wild-type controls. Our results are consistent with prior studies showing that *mitfa*-deficient zebrafish activate distinct signaling pathways, favoring YAP over MAPK signaling, and give rise to tumors with reduced latency compared to wild type tumors.^5^ We observed *yap1* expression in melanocyte progenitor cells of *mitfa*-deficient and wild type zebrafish, suggesting that Yap1 activity may be carried over from the progenitor cell during transformation. While germline *mitfa* loss accelerates tumor onset, our data suggest this effect is not solely due to the deletion of Mitfa from differentiated melanocytes. Instead, we propose that germline *mitfa* deficiency expands a population of *pax3a*-positive melanocyte progenitor cells, which are more susceptible to GNAQ-induced transformation. Alternatively, *mitfa* loss may impair the differentiation of melanocyte progenitors, preventing the formation of GNAQ^Q209L^-positive nevi and promoting direct progression to malignancy. Another explanation for decreased tumor latency in *casper* zebrafish could be that germline loss of *mitfa* has effects on other cell types, such as immune cells, which may influence tumor behavior.

Although Mitfa-independent melanocytic cells are unlikely to be the cellular origin of CM in zebrafish models, studies have shown that MITF-independent BRAF-positive melanoma cells are central to disease recurrence^65^. GNAQ-driven transformation may be facilitated by Mitfa paralogs such as Tfec and Tfeb. The Mitfa paralog Tfec has been shown to activate pigmentation genes in both the retinal pigment epithelium and melanocyte progenitor cells. Tfec can also rescue ectopic melanocytes in *nacre* zebrafish embryos.^42,66^ Moreover, we have recently shown that the MITF paralog TFE3 promotes cellular plasticity in MITF-low melanoma cells,^67^ suggesting that MITF paralogs are compelling candidate transcription factors to drive UM onset in *casper* zebrafish.

In our zebrafish study, melanocyte progenitors inherently express neural crest-associated programs, which are retained during GNAQ^Q209L^-driven transformation. Re-analysis of mouse and human single-cell datasets reveals a subpopulation of *PAX3*-positive melanocytes, suggesting that similar progenitor-like states may be conserved in mammals. Together, these findings highlight how lineage plasticity contribute to UM initiation and progression across species. Such observations are supported by an elegant study by Xu, Karreth and colleagues, who established an immune-competent mouse UM model showing that transcriptional plasticity from melanocytic to neural crest-like states drives tumor progression.^68^

Identifying the transcriptional and signaling mechanisms within *mitfa*-independent melanocyte progenitor cells that facilitate UM onset and progression will have significant clinical implications, particularly for the differential diagnosis of high-risk lesions and the design of targeted therapies.

### Limitation of study

The use of *tp53* and *ptena/b* mutations in our UM model does not fully recapitulate the genetics of human UM. Mutations in *TP53* and *PTEN* are rarely observed in UM, where loss-of-function mutations in *BAP1* represent the most frequent tumor suppressor alteration.^39,69,70^ However, both *TP53* and *PTEN* signaling pathways have been shown to be altered in UM pathogenesis. *MDM2* overexpression, an upstream negative regulator of p53, and downregulation of *PERP*, a downstream effector of p53, have been demonstrated in many UM patients and are associated with worse outcome.^71–75^ Studies have also shown that over half of UM tumors have decreased PTEN immunostaining and its loss is correlated with decreased survival.^39–41^ Whether genetic knockout of these tumor suppressors recapitulate the dysregulation of these pathways in human UM remains an open question. Another limitation to our approach when examining the differences between genetically identical skin and eye-derived tumors is the inability to distinguish between microenvironmental effects and differences in cell of origin (melanocytes in the skin versus in the eye). To address this question, a syngeneic or immunodeficient model could be applied to transplant cells from a primary UM tumor into both anatomical sites.

Additionally, our use of the *mitfa* promoter to drive oncogene expression presents a challenge in defining the exact cell of origin, as this promoter is active in both melanocyte progenitors and differentiated melanocytes. As such, it remains unclear whether tumors in *mitfa* wild-type fish (or those with conditional *mitfa* deletion) arise from a progenitor or a mature melanocyte. Lineage tracing studies will be essential to determine whether differentiated melanocytes possess the capacity to initiate primary UM in *mitfa*-competent or conditional *mitfa*-KO zebrafish.

## Acknowledgments

This work was supported by the Holden Comprehensive Cancer Center (P30 CA086862). Additional support to C.K. was provided by an institutional research grant from the American Cancer Society administered through the HCCC (IRG-18-164-43), the Melanoma Research Alliance (ID#1426690) and Outrun the Sun in honor of Doug Stickney. J.Y. and D.R. were supported by NIH predoctoral fellowships (T32 GM144636 and T90 DE023520, respectively). Additional NIH support was provided to C.K. (R03 CA297549-01), R.A.C. (R01 AR062457), R.F.M. (P30 EY025580), and D.L. (R03 CA288281-01). Further funding was provided by the Leo Foundation (LF-OC-21-000888) to D.L., the Wisconsin Partnership Program (AAN4246) to A.B., and the U.S. Department of Defense (ME240045) to A.B. The funders had no role in study design; data collection, analysis, or interpretation; manuscript preparation; or the decision to publish.

---

**FIGURE S1.**
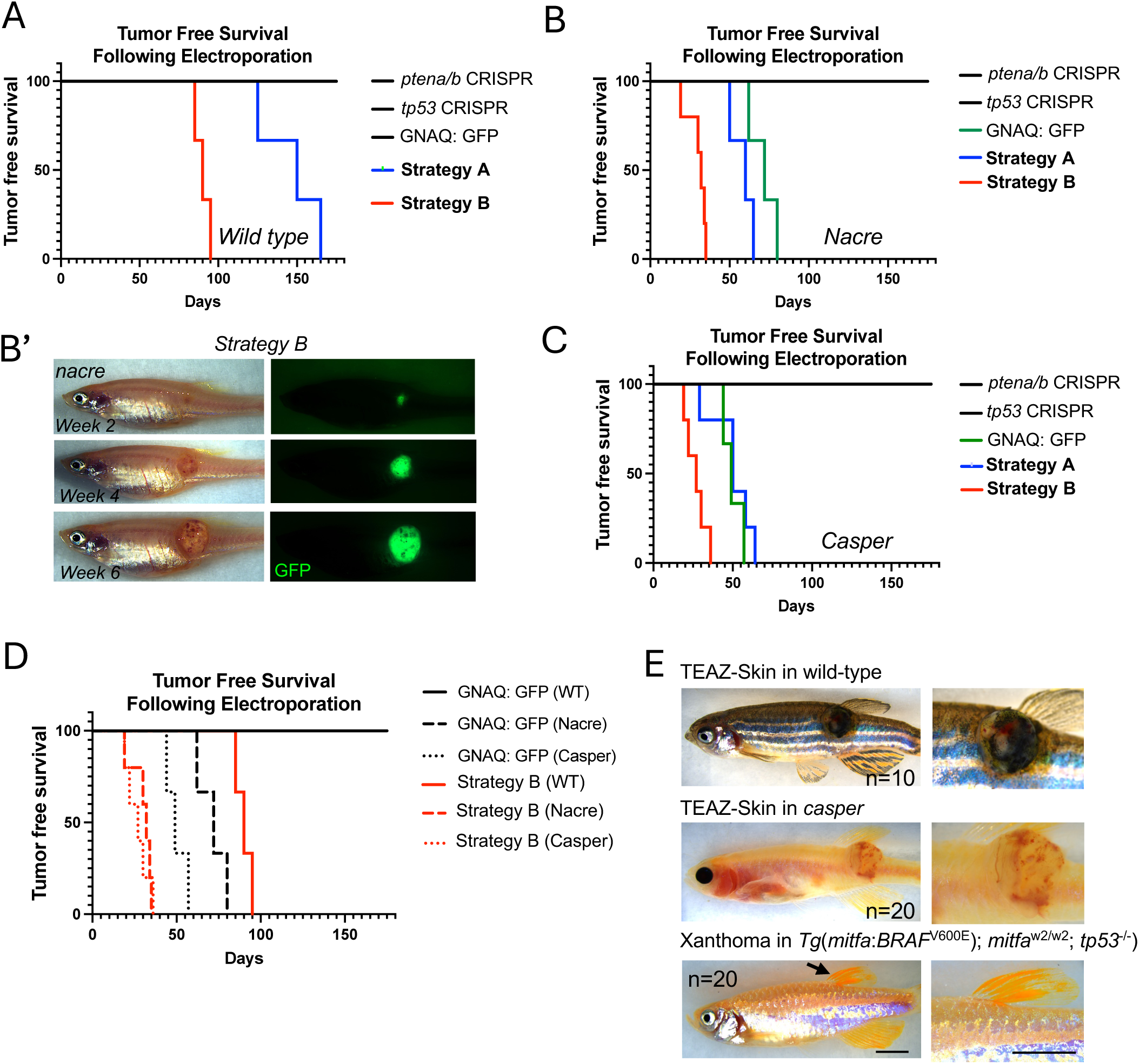

**FIGURE S2.**
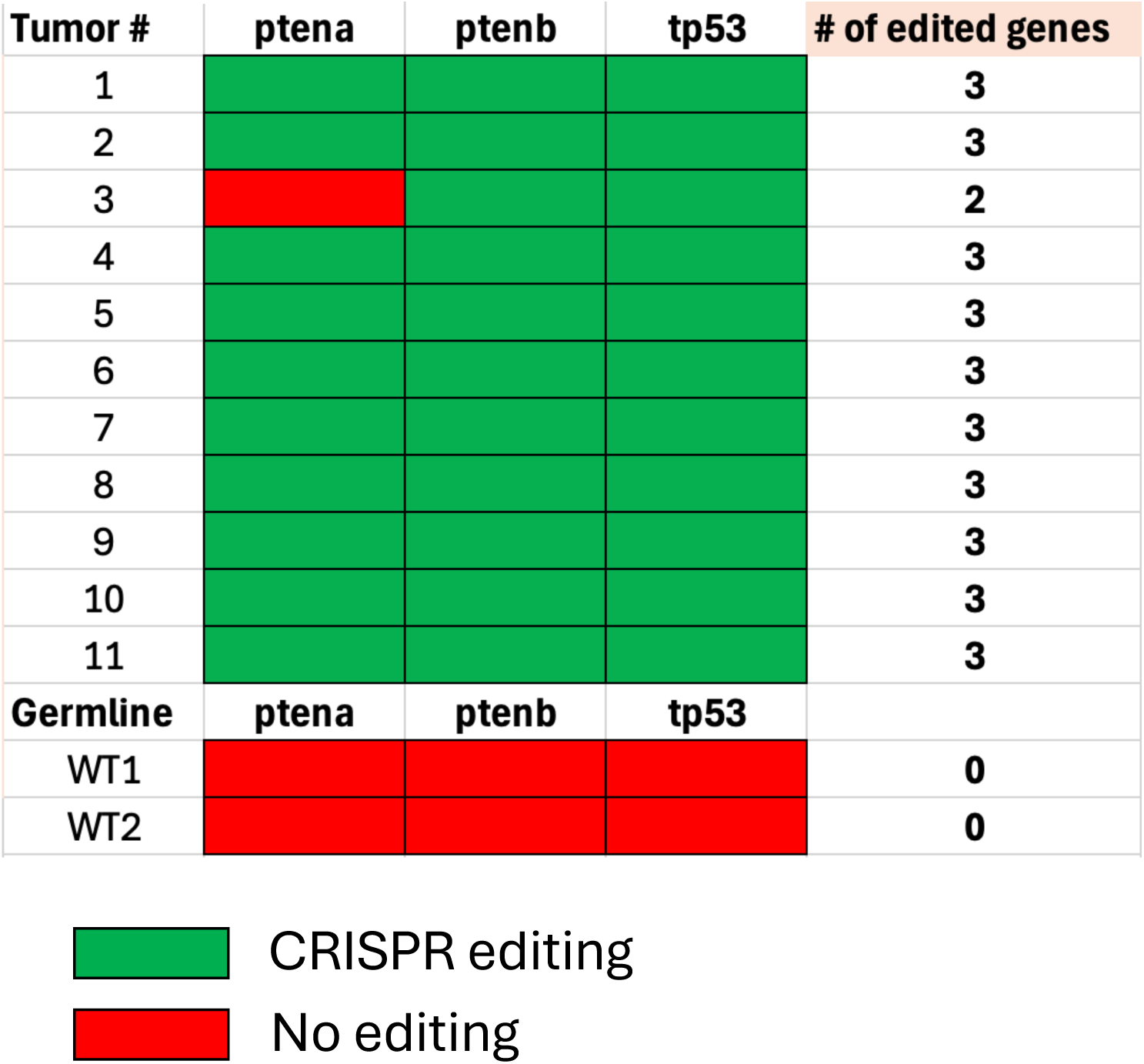

**FIGURE S3.**
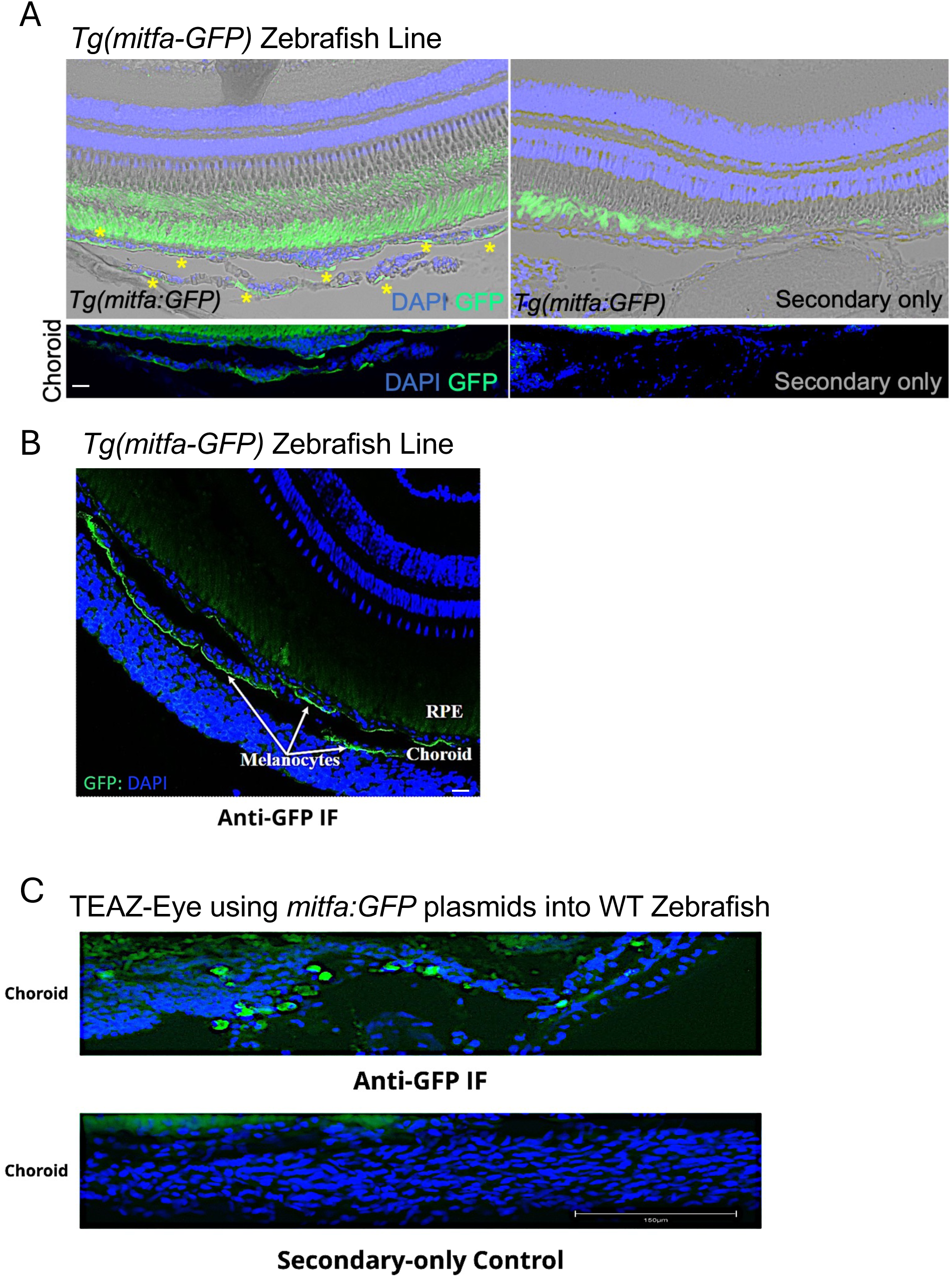

**FIGURE S4.**
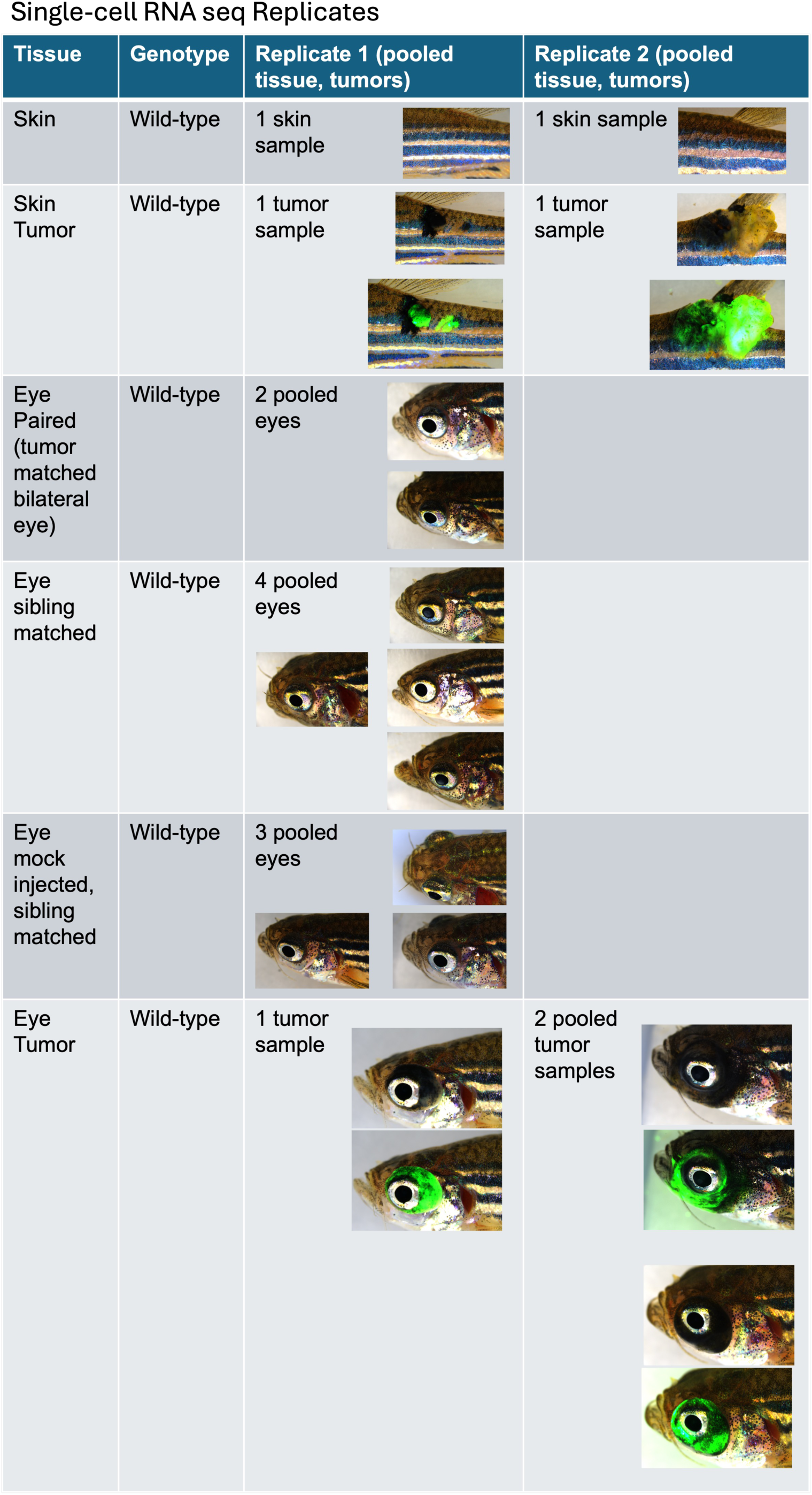

**FIGURE S5.**
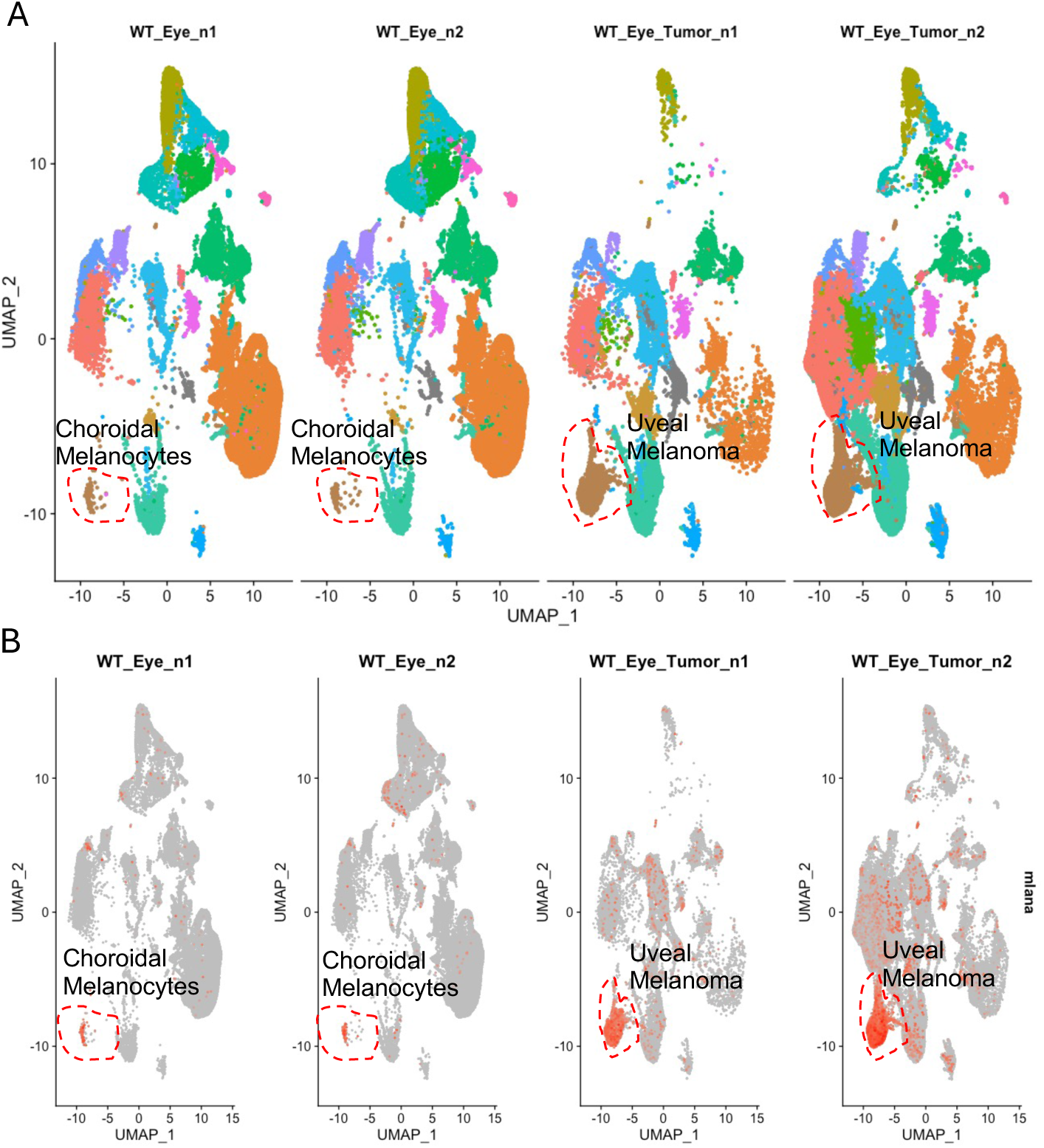

**FIGURE S6.**
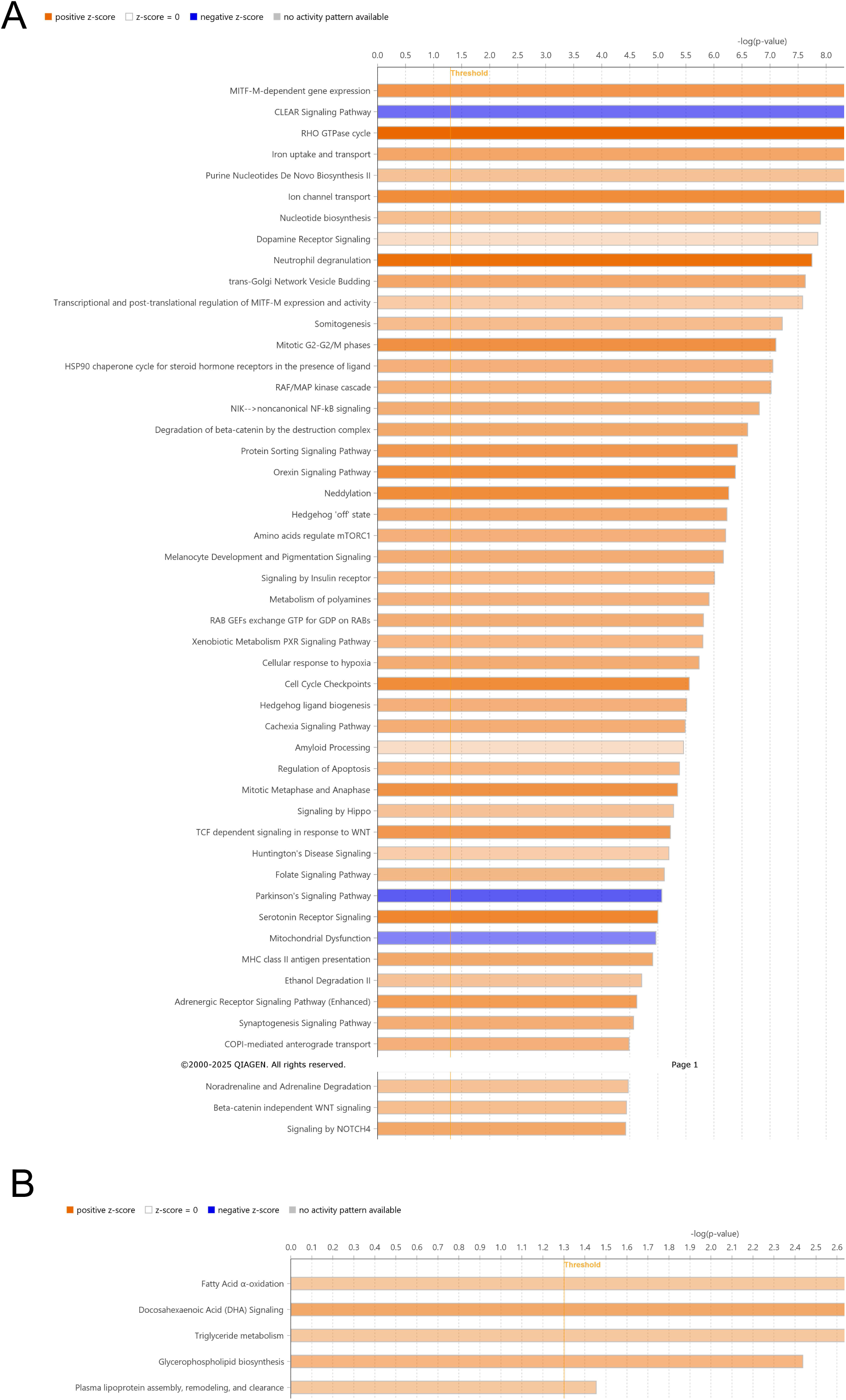

**FIGURE S7.**
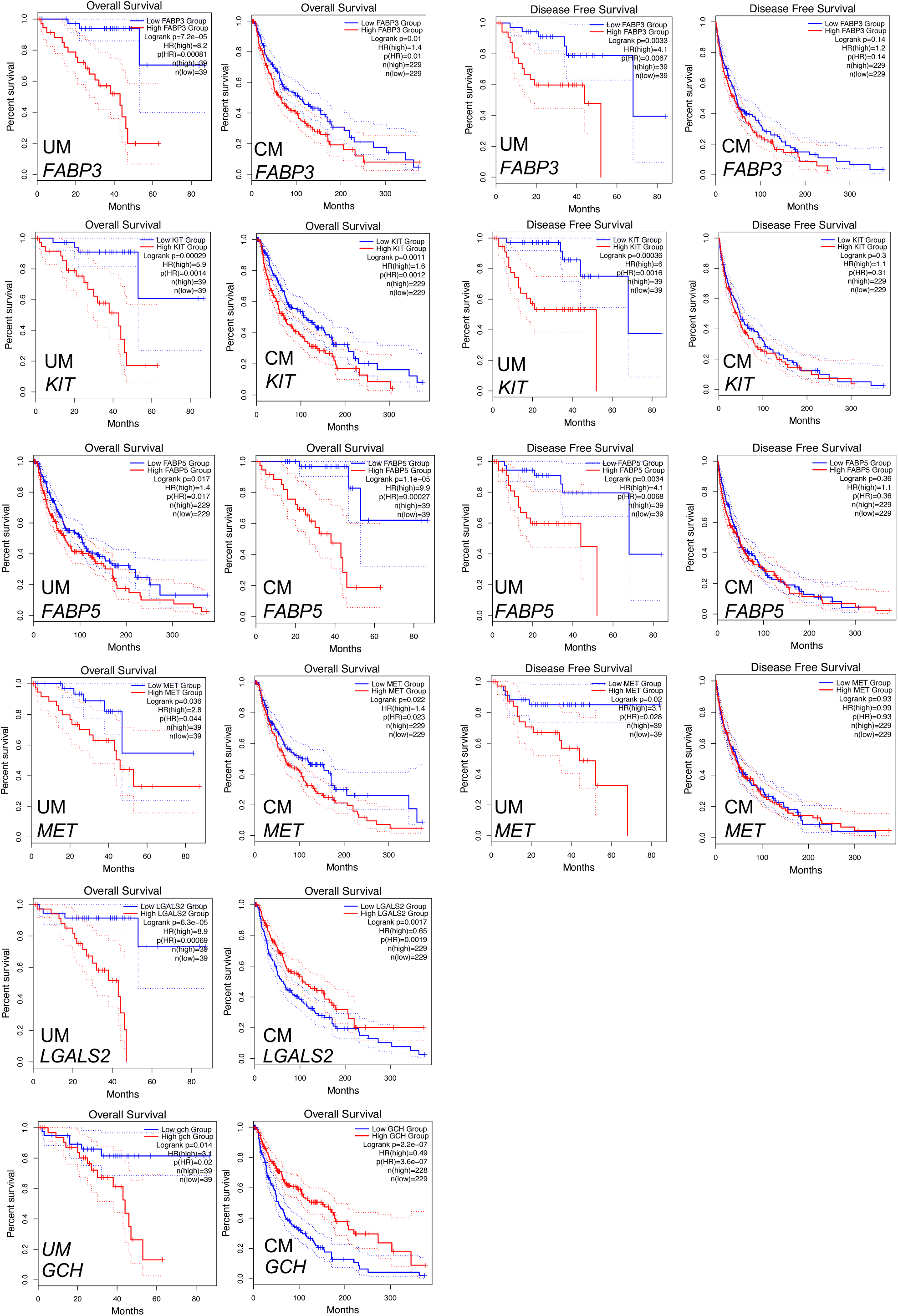

**FIGURE S8.**
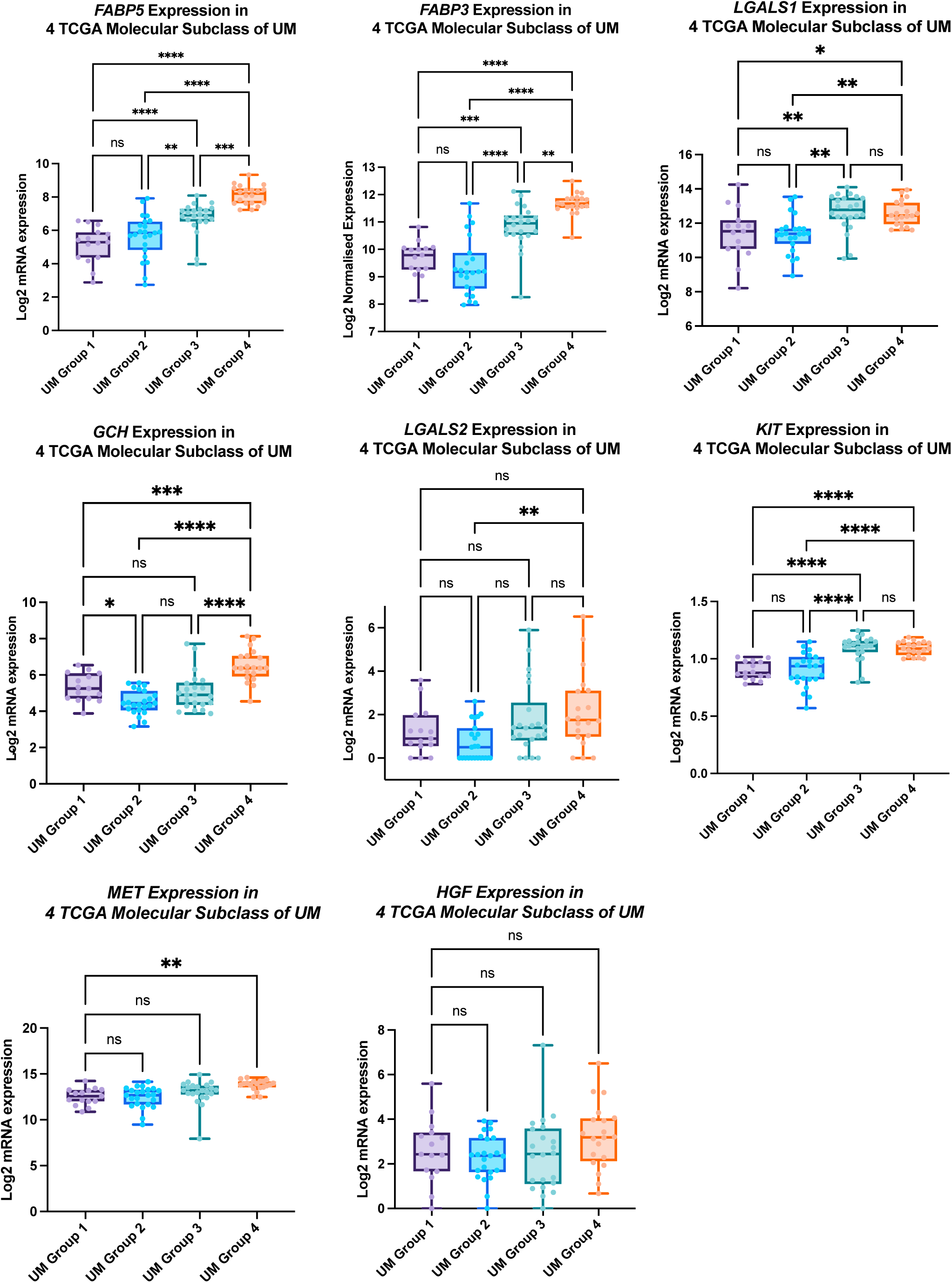

**FIGURE S9.**
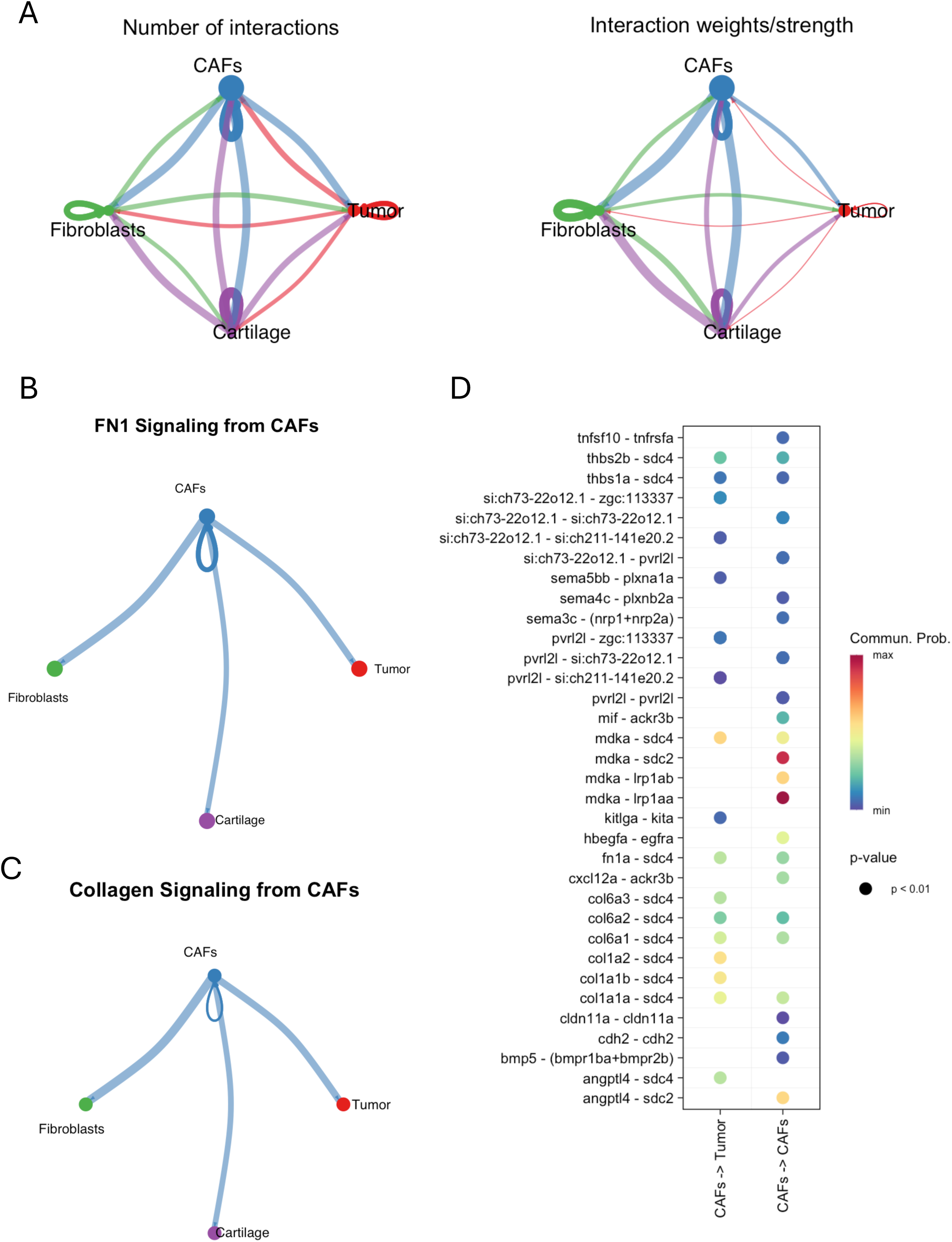

**FIGURE S10.**
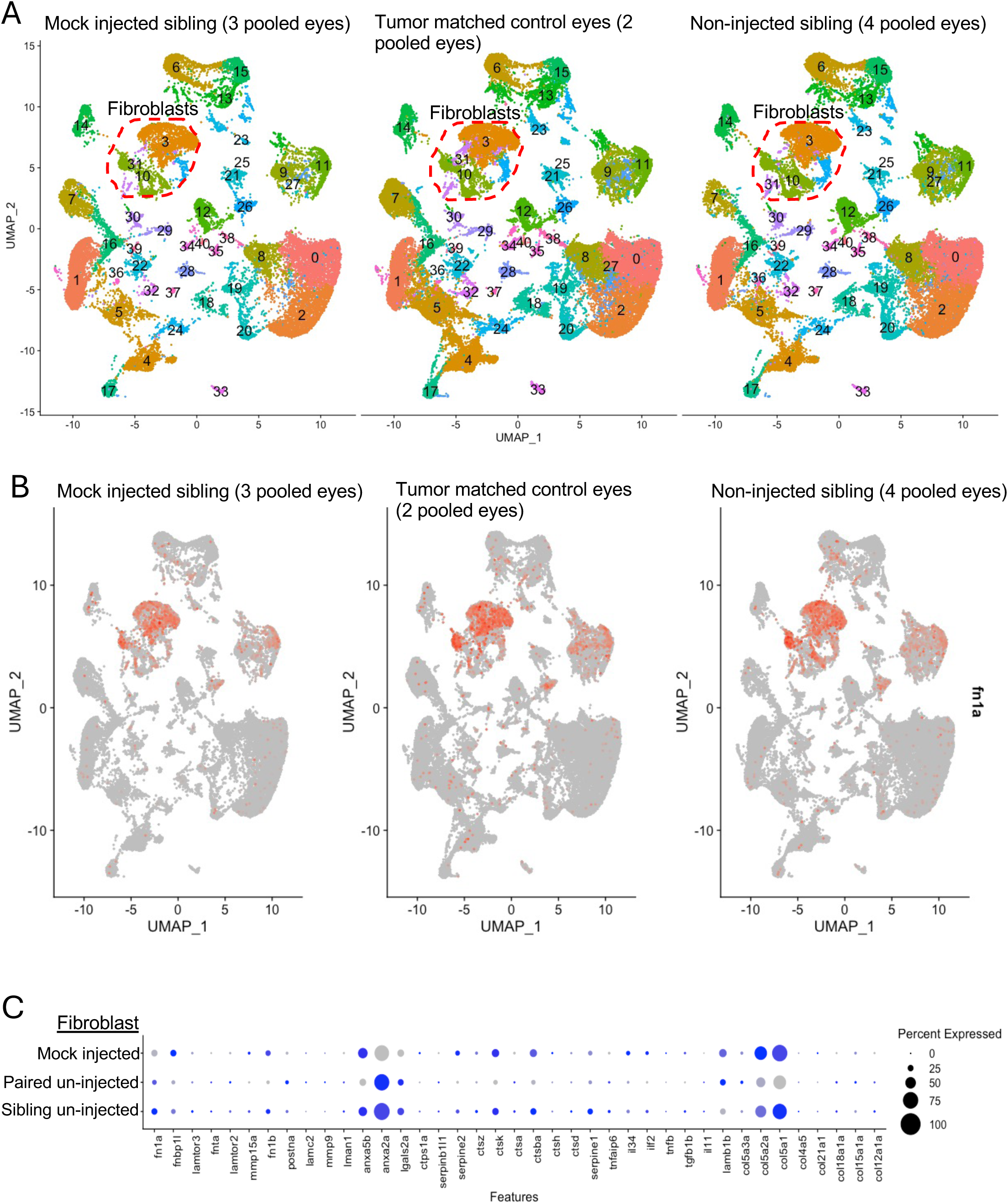

**FIGURE S11.**
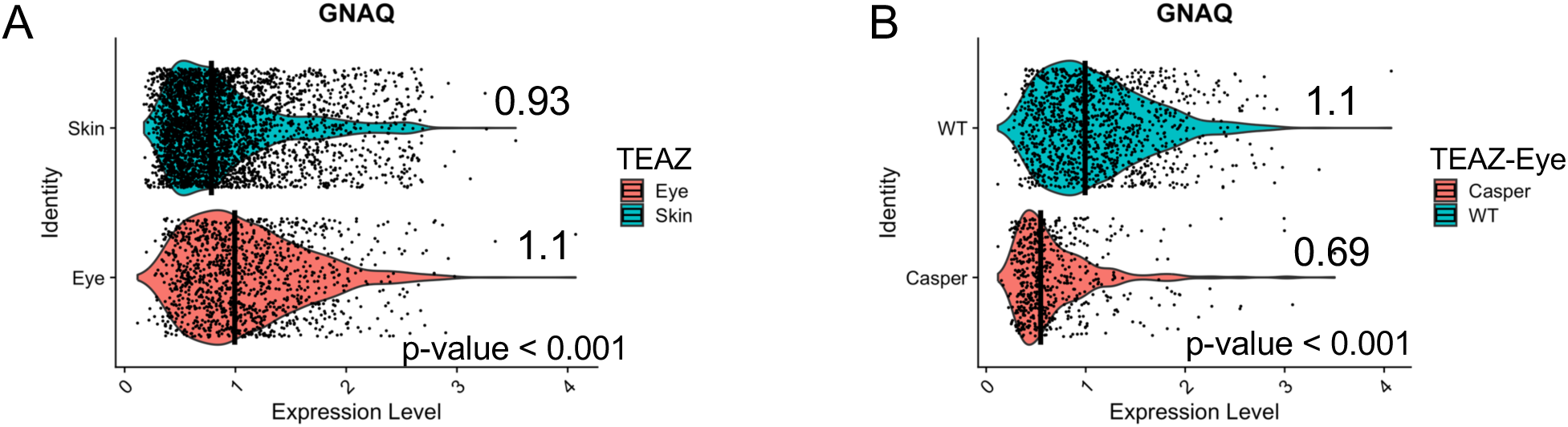

**FIGURE S12.**
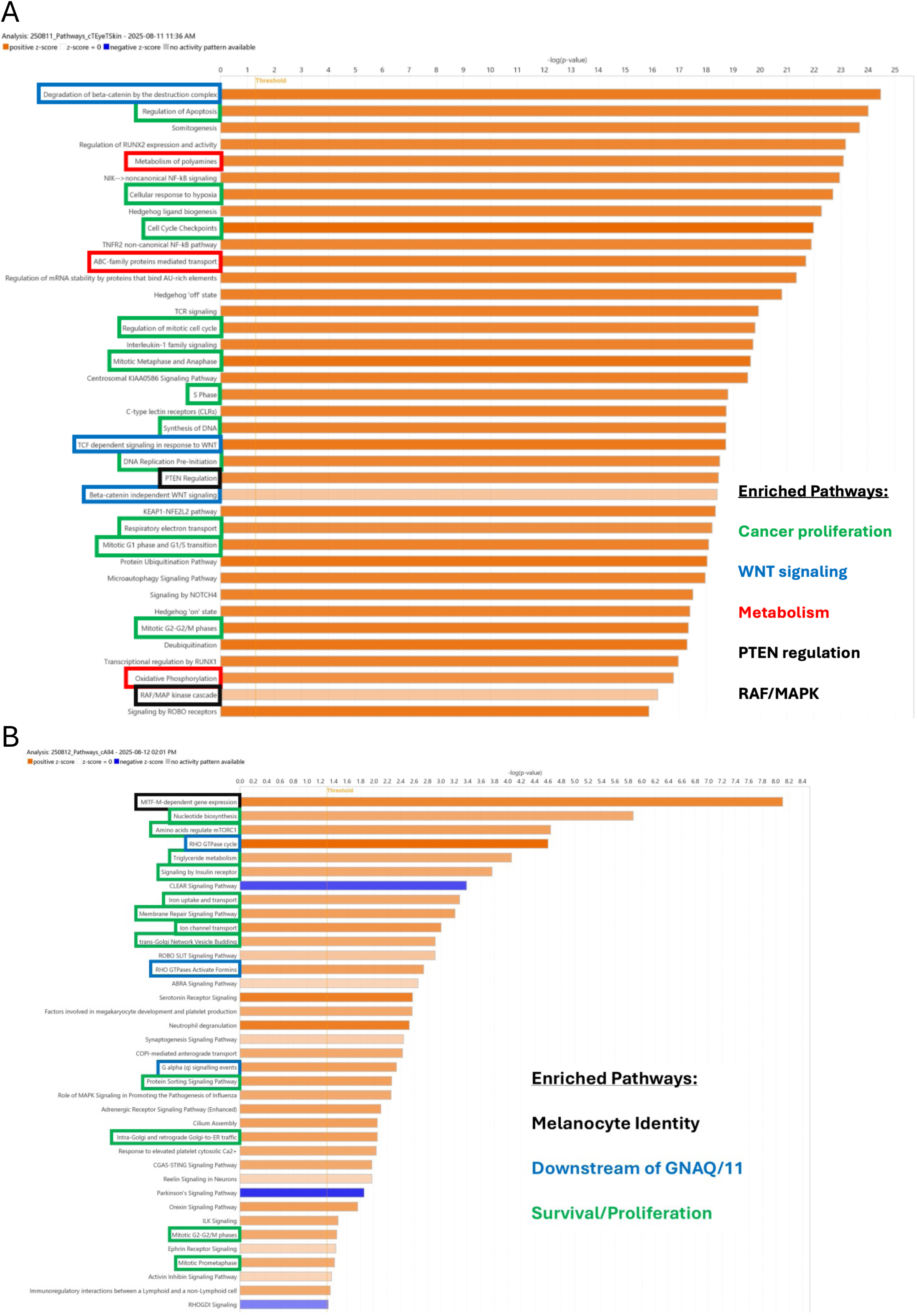

**FIGURE S13.**
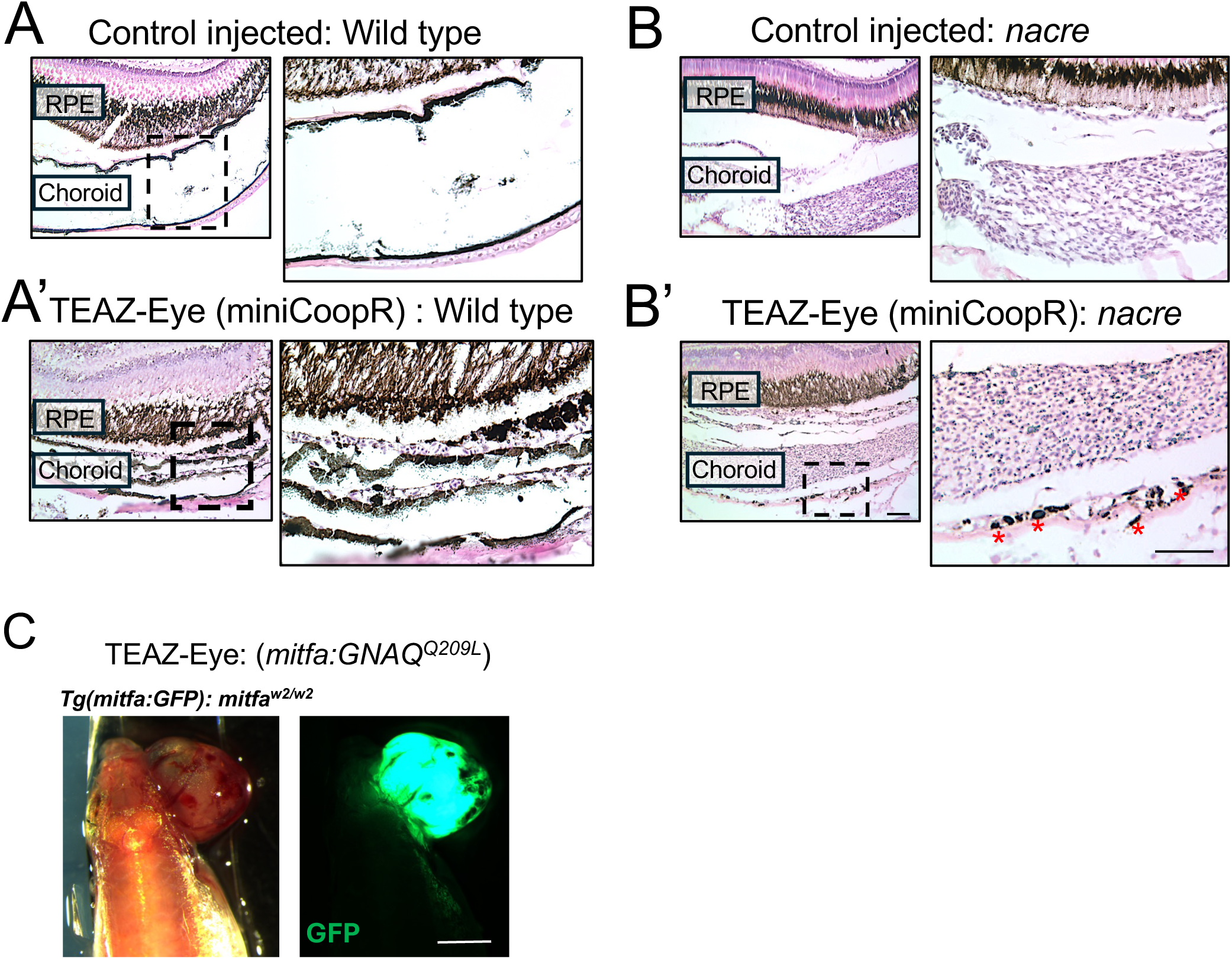

**FIGURE S14.**
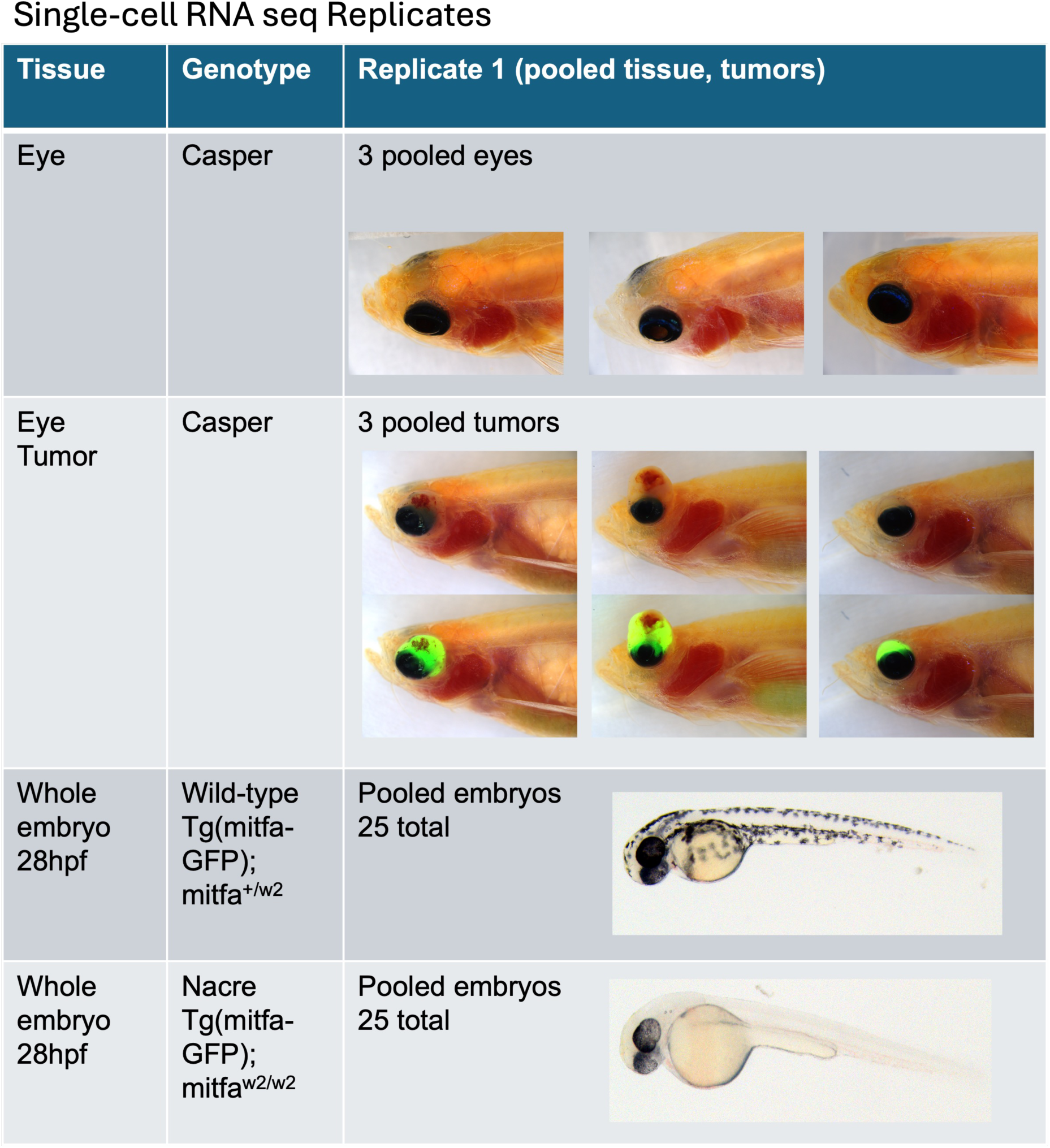

**FIGURE S15.**
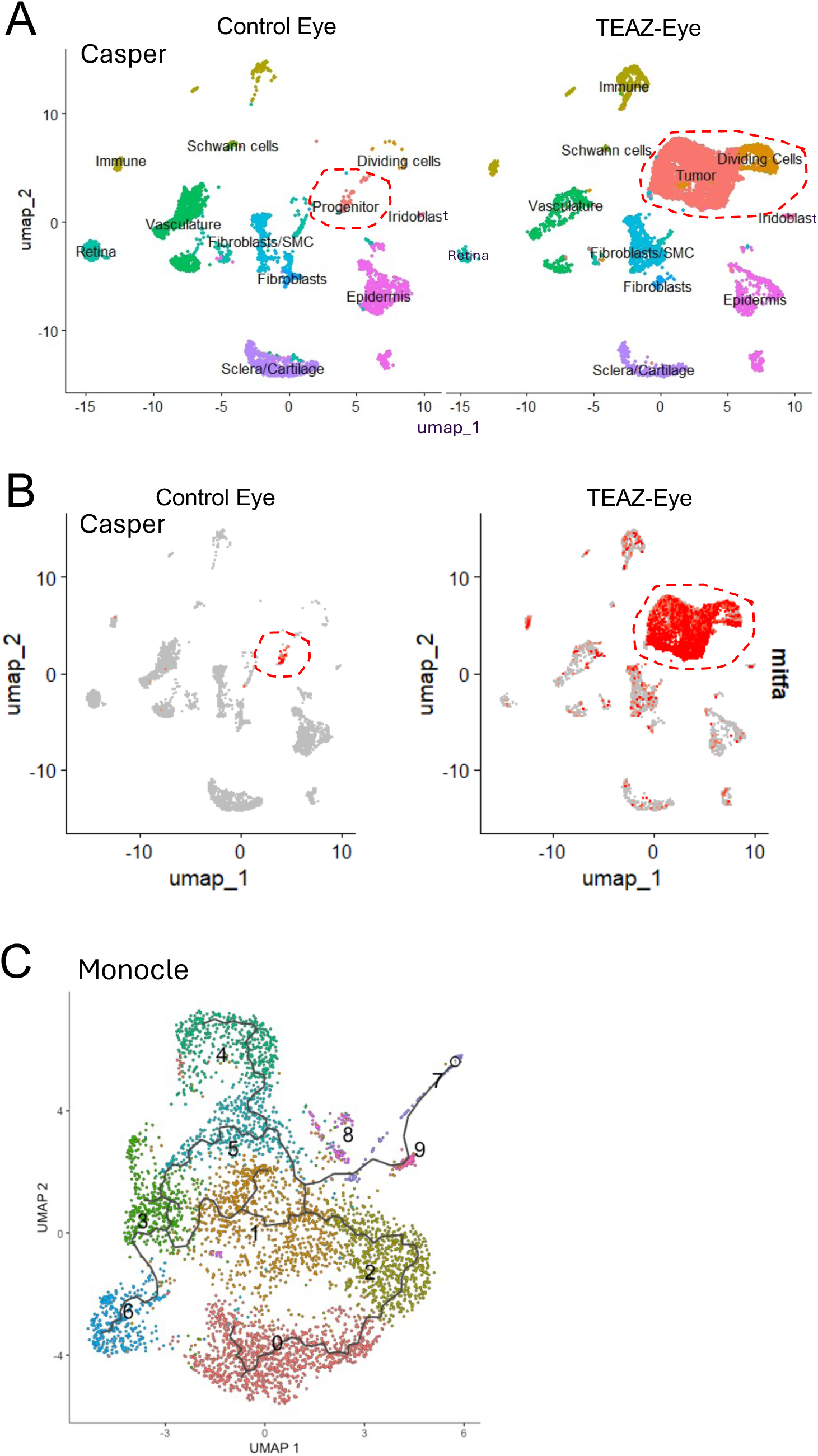

**FIGURE S16.**
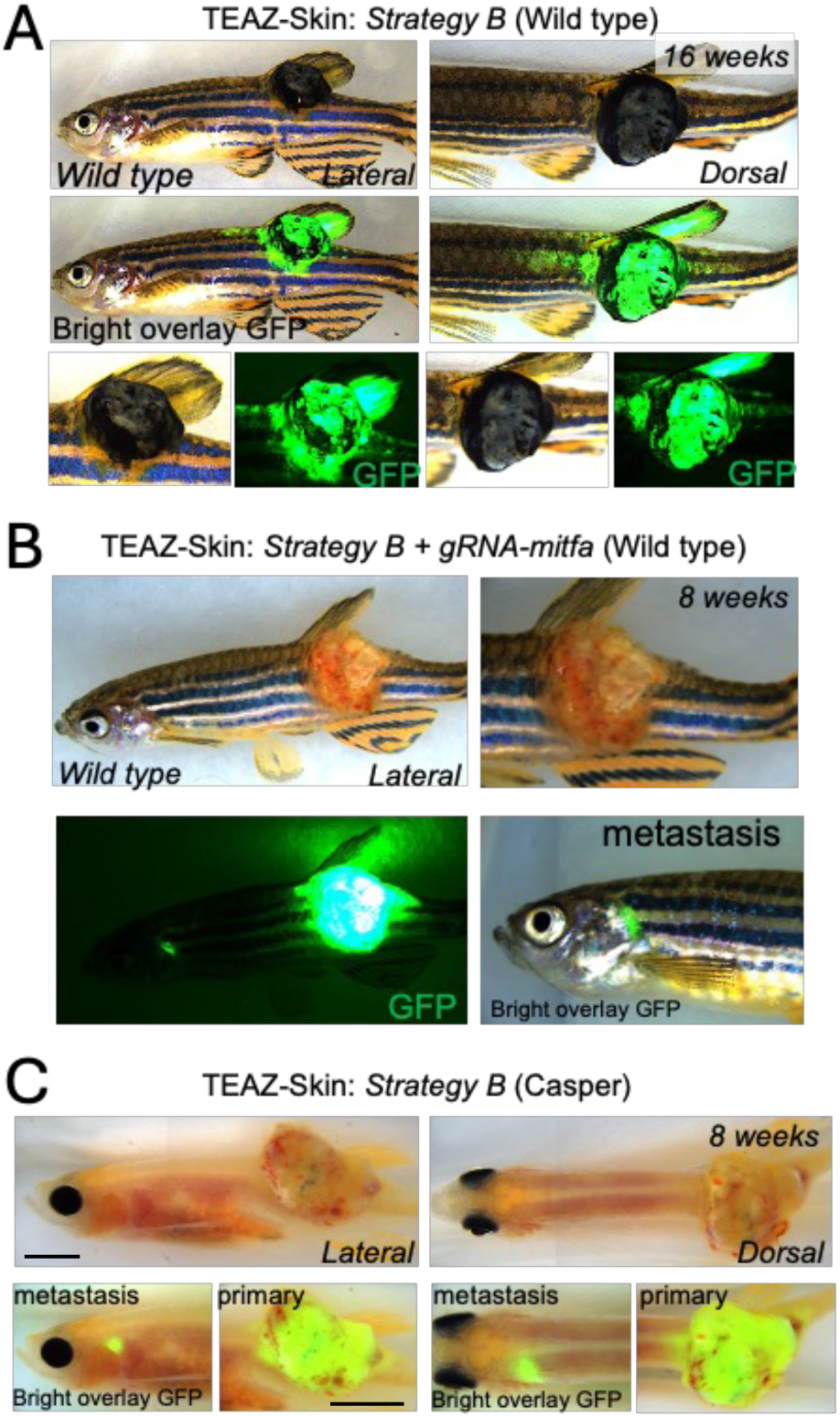

**FIGURE S17.**
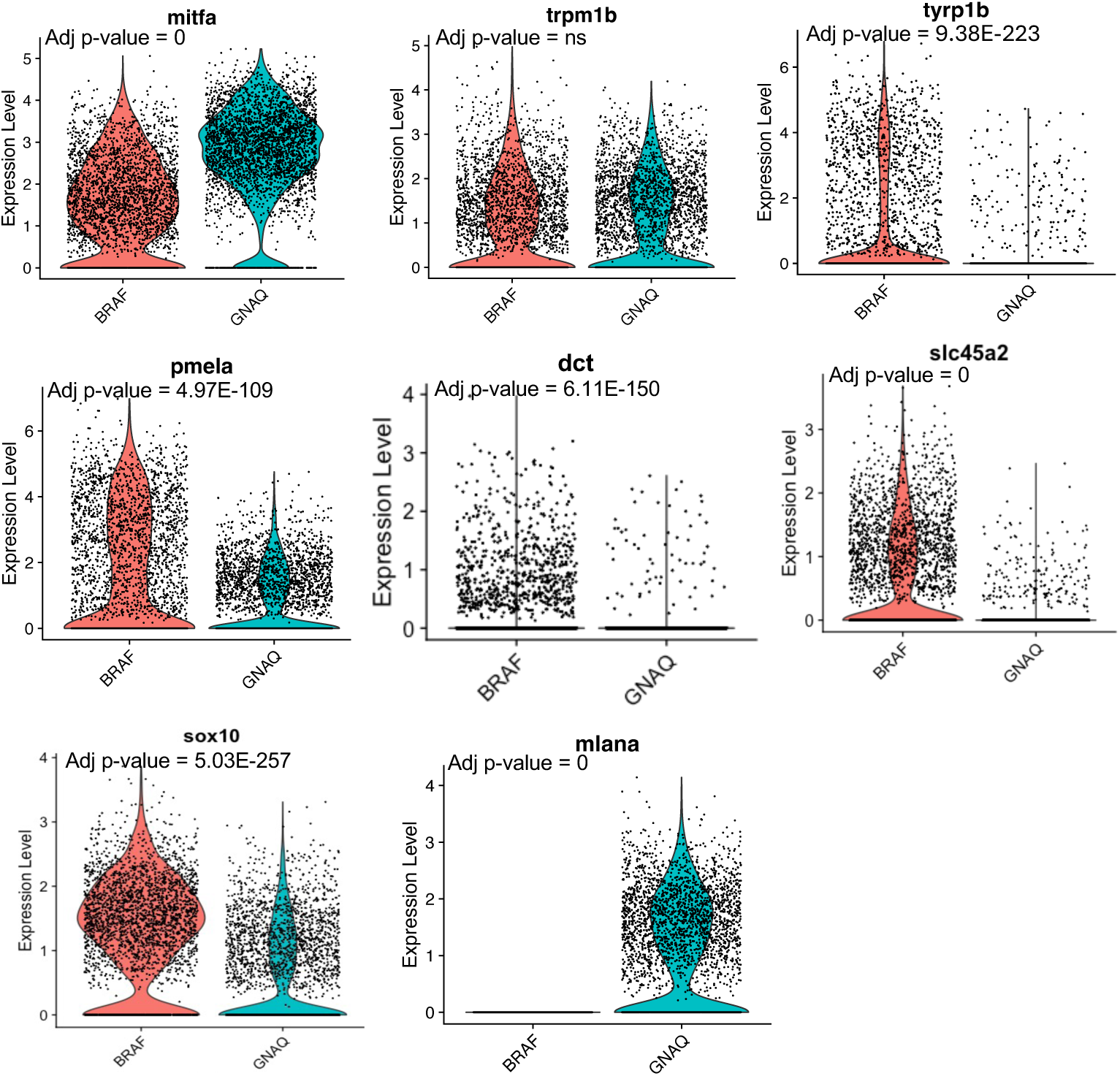

**FIGURE S18.**
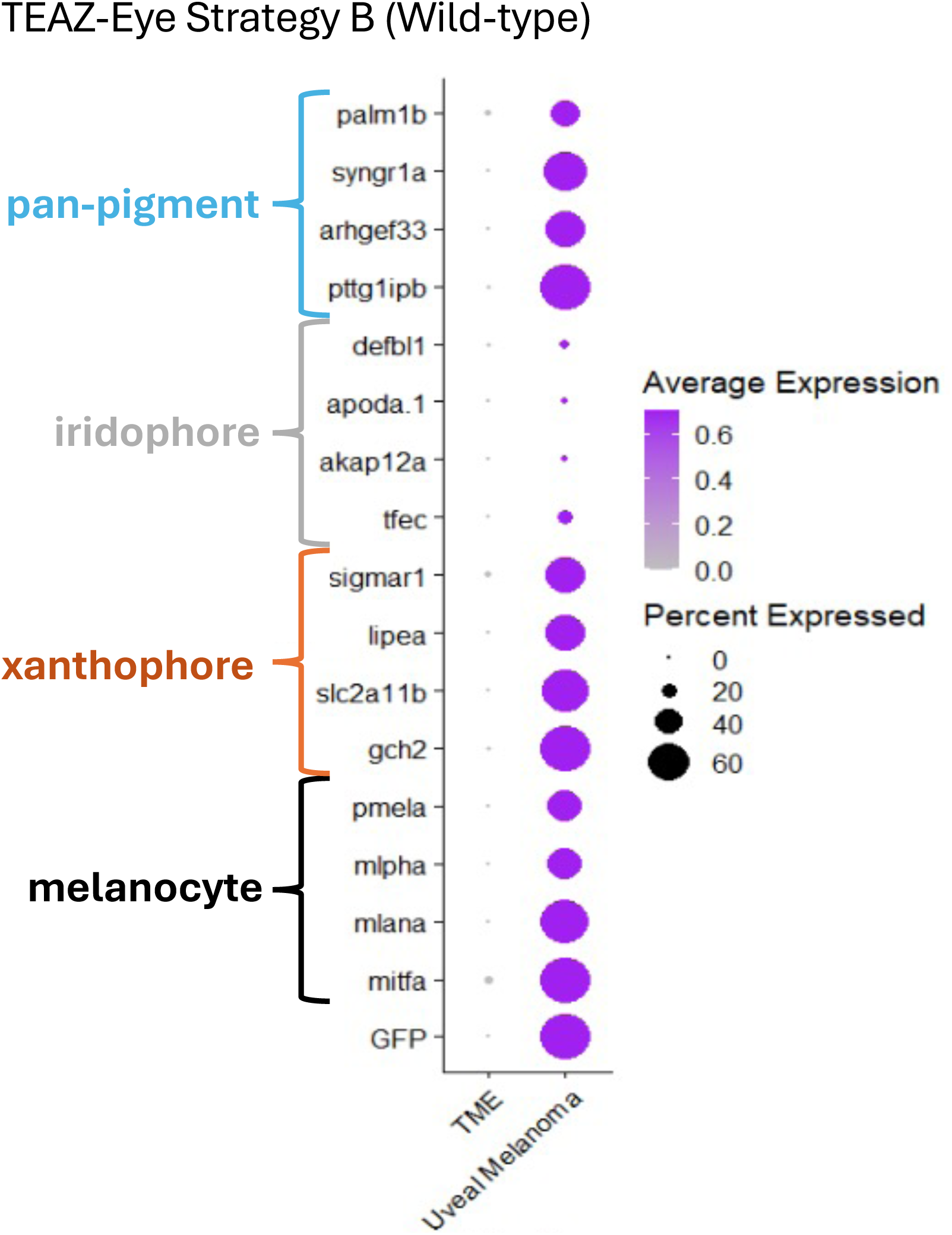

**FIGURE S19.**
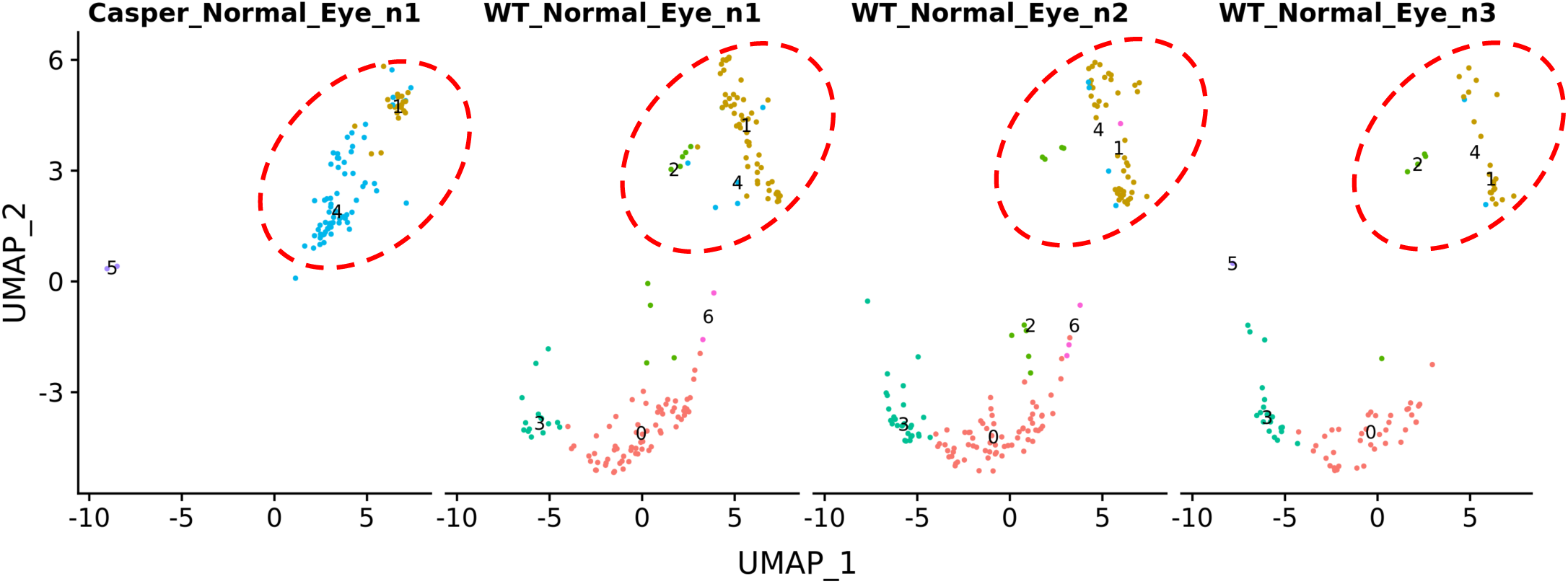

**FIGURE S20.**
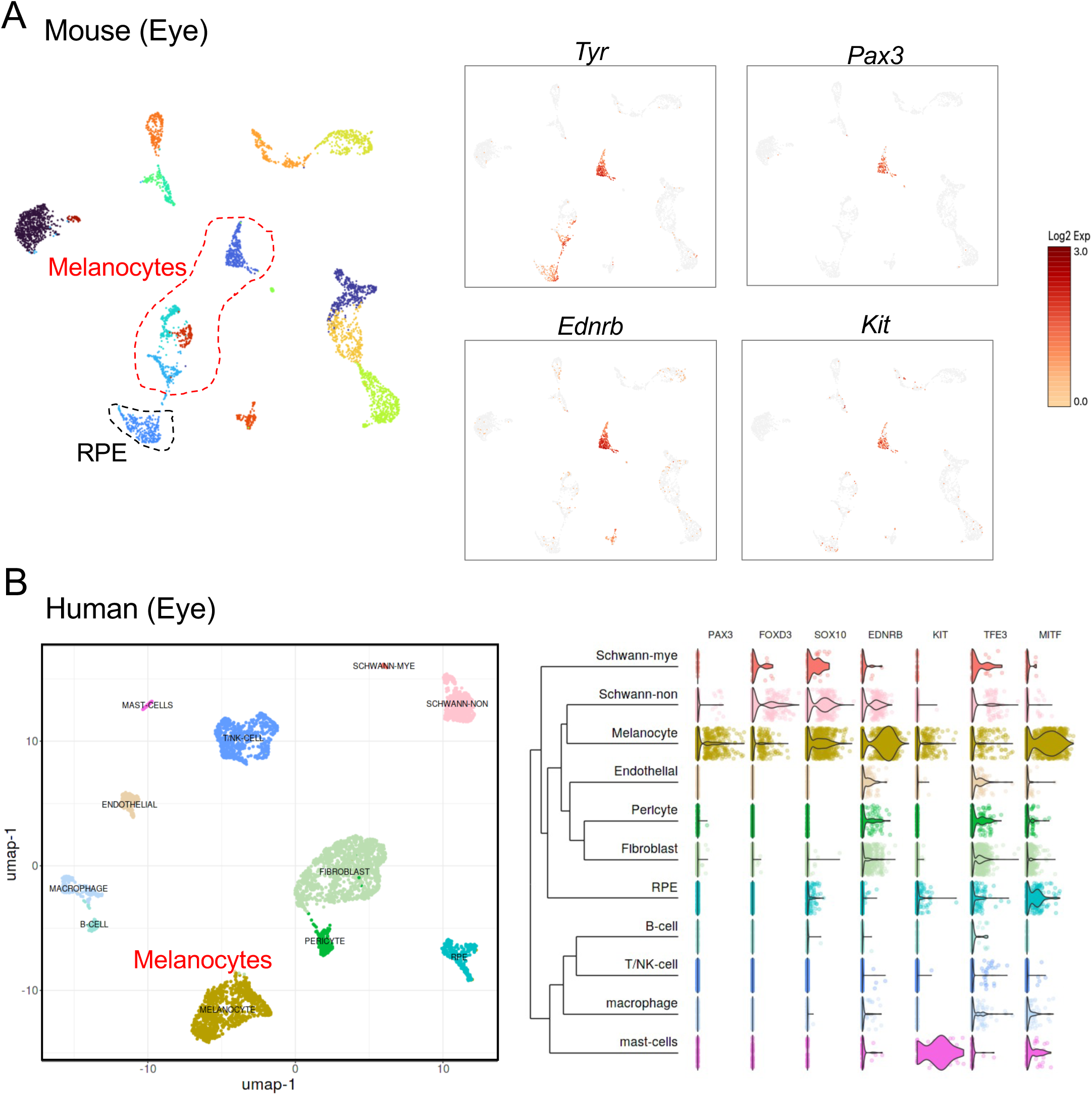

**FIGURE S21.**
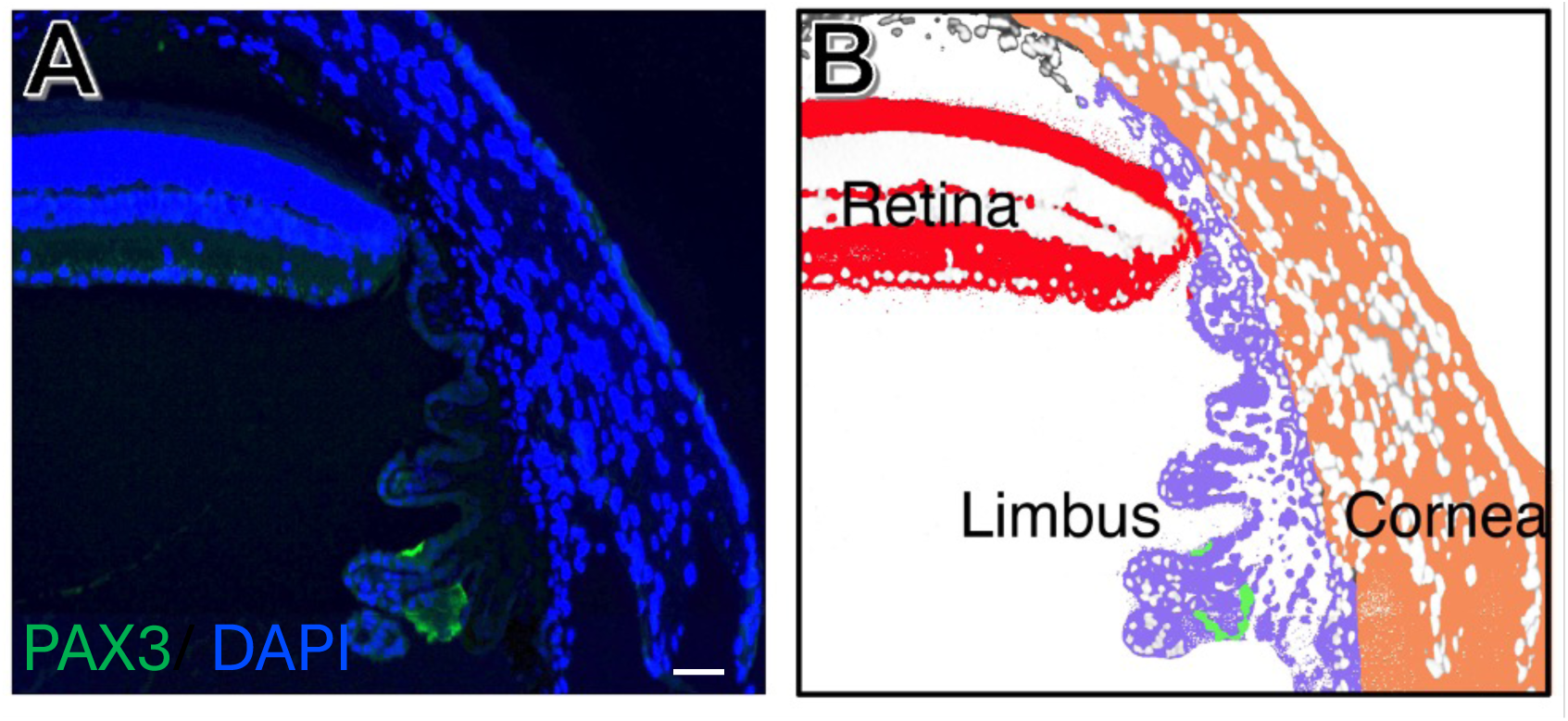

