## Supplementary material for "GNAQ Induces Melanomagenesis in Mitfa-Independent Melanocyte Progenitors in a Zebrafish Model of Uveal Melanoma": Figures_S1-21

**Figure S1: Mitfa-deficient zebrafish have decreased tumor latency compared to wild-types by TEAZ-Skin**.

(**A**) Kaplan-Meier curves estimating tumor-free survival, defined as the time from electroporation until the first visible appearance of GFP+ nodular tumors, in adult wild-type zebrafish via TEAZ-Skin. Injections contained *mitfa*:Cas9 with *U6*:gRNA-*ptena* + *U6*:gRNA-*ptenb* (*ptena/b* CRISPR), *mitfa*:Cas9 with *U6*:gRNA-*tp53* (*tp53* CRISPR), *mitfa*:*GNAQ*^Q209L^-*mitfa*:GFP (GNAQ: GFP), strategy A plasmids, or strategy B plasmids (n=3 per condition). (**B**) Kaplan-Meier curves estimating tumor-free survival in adult *nacre* (*mitfa*^w2/w2^) zebrafish via TEAZ-Skin. Same injection mixes as in (A). (n=5 for Strategy B, n=3 for rest) (B’) Representative brightfield and fluorescent images shown at 2, 4, and 6 weeks for strategy B injection in *nacre*. (**C**) Kaplan-Meier curves estimating tumor-free survival in adult *casper* (*mitfa*^w2/w2^, *mpv17*^a9/a9^) zebrafish via TEAZ-Skin. Same injection mixes as in (A). (n=5 for Strategies A and B, n=3 for rest) (**D**) Kaplan-Meier curves estimating tumor-free survival in adult wild-type, *nacre*, and *casper* zebrafish via TEAZ-Skin. Oncogene alone (GNAQ: GFP) is compared to strategy B plasmids for each genotype. (n=5 for strategy B in *nacre* and *casper*, n=3 for rest). (**E**) Representative images of Strategy B TEAZ-Skin in wild-type and *casper* and a xanthoma that formed in a *Tg(mitfa:BRAF^V600E^); mitfa^w2/w2^; tp53^-/-^* zebrafish. Note the bright orange color of the xanthoma, indicated by the black arrow. Scale bars = 0.5 cm

**Figure S2:** **Targeted Sanger sequencing to confirm CRISPR gene editing following TEAZ-Skin injection using Strategy B plasmids.** 11 tumors and 2 wild-type tissue controls were used for INDEL analysis to verify gene editing in *tp53, ptena,* and *ptenb.* Green box indicates successful gene editing; red box indicates no gene editing. Total number of edited genes per sample are indicated in the right-most column.

**Figure S3: Immunofluorescence of the choroid using an anti-GFP antibody**

(**A**) *Tg*(*mitfa*:GFP) with 10x magnification and in (**B**) *Tg*(*mitfa*:GFP) following quenching of autofluorescence. Scale bar = 20μm (**C**) TEAZ-Eye–injected zebrafish expressing a GFP construct driven by the *mitfa* promoter (*mitfa:GFP*) with 40x magnification. Sections were counterstained with DAPI to visualize nuclei. Scale bar = 150μm

**Figure S4: Gross images of tissue and tumors used for single cell RNA-Seq in wild-type zebrafish.**

**Figure S5: UMAP of normal eyes and TEAZ-Eye uveal melanoma in wild-type zebrafish (Strategy B).** (**A**) Integrated object is split by sequencing replicates (top) and feature plots showing *mlana* RNA expression to identify normal choroidal melanocytes and uveal melanoma cells (bottom). Color intensity (grey to red) reflects normalized average expression levels (low to high). The red dotted line indicates the melanocyte and uveal melanoma clusters.

**Figure S6: Ingenuity Pathway Analysis of genes enriched in TEAZ-Eye compared to the tumor microenvironment**. (**A**) The top 49 pathways enriched in TEAZ-Eye. (**B**) Select pathways enriched in TEAZ-Eye pertaining to fatty acid oxidation and signaling. Color intensity indicates the sign and magnitude of the z-score from blue (negative) to orange (positive). Significance is indicated by the -log(p-value).

**Figure S7:** **Overall and disease-free survival analysis:** Overall survival of patients with uveal melanoma (UM) and cutaneous melanoma (CM) based on expression levels of *FABP3*, *KIT, FABP5, MET, LGALS2,* and *GCH*. Disease-free survival in UM and CM patients based on *FABP3, KIT, FABP5* and *MET* expression. Dotted lines represent 95% confidence intervals. Data derived from TCGA datasets.

**Figure S8: TCGA group analysis:** Normalized RNA expression of *FABP5, FABP3, LGALS1, GCH, LGALS2, KIT, MET* and HGF across the 4 TCGA molecular subclasses. ^55^ Groups were compared by ANOVA with post-hoc Tukey test. ns = not significant, * p-value<0.05; ** p-value<0.01; ***p-value<0.001; ****p-value<0.0001.

**Figure S9: CellChat analysis of intercellular communication between zebrafish fibroblast, CAF, and UM cell clusters.** Cell-cell communication analysis was performed using CellChat on fibroblast, CAF, cartilage, and UM tumor cell clusters defined in Figure 2. (**A**) Circle plots showing the number of inferred interactions (left) and interaction strength/weight (right) among the indicated cell populations. Edge thickness is proportional to the number or strength of predicted ligand-receptor interactions. (**B**) FN1 signaling network highlighting CAFs as the dominant source of FN1 pathway signaling to UM tumor cells and other stromal populations. (**C**) Collagen signaling network demonstrating CAF-derived collagen pathway interactions with UM tumor cells. (**D**) Dot plot of ligand-receptor pairs contributing to CAF-tumor communication. Dot color indicates communication probability (blue to red), and dot size reflects statistical significance (p < 0.01).

**Figure S10: scRNA-Seq on control wild-type zebrafish eyes.** Mock injected wild-type zebrafish eyes do not show an increase in activated fibroblast markers as compared to uninjected controls. (**A**) UMAP representation of wild-type normal eyes from three different sources: mock injected, sibling-matched (3 pooled eyes), uninjected tumor-paired (2 pooled eyes), and uninjected sibling-matched (4 pooled eyes). The red dotted line indicates the fibroblast clusters. (**B**) Feature plots showing *fn1a* RNA expression across wild-type eye control groups. Color intensity (grey to red) reflects normalized average expression levels (low to high). (**C**) Dot plot of fibroblast activation marker genes in the fibroblast clusters of wild-type eye control groups. Dot size indicates the percentage of cells expressing each gene; color intensity (grey to blue) reflects normalized average expression levels (low to high).

**Figure S11: Oncogenic GNAQ expression in wild-type and casper eye tumors.** Violin plots showing GNAQ^Q209L^ expression in tumor cell clusters from (**A**) wild-type TEAZ-Eye and TEAZ-Skin tumors and (**B**) wild-type and *casper* TEAZ-Eye tumors. Statistical significance was assessed using the Wilcoxon test on scRNA-Seq profiles.

**Figure S12: Ingenuity Pathway Analysis of enriched genes in TEAZ-Eye and TEAZ-Skin compared to the combined tumor microenvironment (eye and skin).** (**A**) Enriched pathways from genes that were common to only TEAZ-Eye and TEAZ-Skin compared to the combined tumor microenvironment. Families of pathways have been identified with a colored box and colored label including: cancer proliferation (green), WNT signaling (blue), metabolism (red), PTEN regulation (black), and RAF/MAPK signaling (black). (**B**) Enriched pathways from genes that were common to all four gene subsets (TEAZ-Eye, TEAZ-Skin, choroidal melanocytes, skin melanocytes) versus their respective combined microenvironments (normal or tumor). Families of pathways have been identified with a colored box and colored label including: melanocyte identity (black), signals downstream of GNAQ/GNA11 (blue), and survival/proliferation (green). Color intensity indicates the sign and magnitude of the z-score from blue (negative) to orange (positive). Significance is indicated by the -log(p-value).

**Figure S13: Melanocyte rescue experiments in *nacre* zebrafish by TEAZ-Eye using plasmids containing *mitfa*:*mitfa*; *mitfa*:GFP (miniCoopR).** (**A-A’**) H&E images (10x and 40x magnification) of wild-type (*AB*) eyes following TEAZ-Eye injection with (**A**) control *mitfa*-GFP or (**A’**) miniCoopR plasmids. (**B-B’**) H&E images (10x and 40x magnification) of *mitfa^w2/w2^* (*nacre*) eyes following TEAZ-Eye injection with (**B**) control *mitfa*-GFP or (**B’**) miniCoopR plasmids, rescued melanocytes in the choroid are indicated by a red asterisk. Scale bar = 50μm. (**C**) Representative dorsal image of a transgenic *Tg(mitfa:GFP); mitfa^w2/w2^* injected with *mitfa*:*GNAQ^Q209L^* following TEAZ-Eye injection. Scale bar = 0.5 cm.

**Figure S14: Sample overview of adult *casper* zebrafish and *nacre* embryos:** Gross images of tissue and tumors used for single cell RNA-Seq in *casper* zebrafish and transgenic *Tg*(*mitfa*:GFP); *mitfa*^+/w2^ and *Tg*(*mitfa*:GFP); *mitfa*^w2/w2^ whole embryos.

**Figure S15: UMAP representation of UM tumors derived from *casper* zebrafish, with accompanying pseudotime analysis.** (**A**) UMAP representation following scRNA-Seq on normal *casper* control eyes (3 pooled eyes) and UM tumors (3 pooled tumors) derived in *casper* (TEAZ-Eye) (**B**) Feature plot showing *GFP* expression and *mitfa*^w2/w2^ expression (**C**) UMAP following re-clustering of *mitfa*^w2/w2^ positive cells from (B), with overlayed unsupervised pseudotime analysis using Monocle analysis.

**Figure S16: Representative images of TEAZ-Skin induced tumors (*Strategy B*)**: (**A**) wild-type at 16 weeks, (**B**) conditional *mitfa*-KO (wild-type) at 8 weeks and (**C**) *casper* zebrafish at 8 weeks post TEAZ-Skin. Brightfield and GFP-overlaid images are shown to highlight primary and metastatic cells in (B-C), and the absence of metastasis in (A). Scale bar = 0.5 cm

**Figure S17:** Violin plots representing *mitfa* expression as well as pigmentation genes in BRAF^V600E^- and GNAQ^Q209L^-positive tumors by single cell RNA-Seq analysis.

**Figure S18:** Dot plot of genes enriched in chromatoblasts and in uveal melanoma tumors derived in wild-type zebrafish following TEAZ-Eye compared to the tumor microenvironment (TME). Genes enriched in a particular pigment cell type are labelled, including: iridophore, xanthophore, and melanocyte.

**Figure S19:** UMAP representation of melanocytes and melanocyte progenitors in *casper* normal eyes and in the three sequencing replicates of wild-type normal eyes. The red dotted line indicates melanocyte progenitors in the *casper* sample and melanocytes that are clustering closely with such progenitors in wild-type samples.

**Figure S20: PAX3 positive melanocytes are observed in mouse and human eyes by scRNA-Seq analysis.** (**A**) UMAP representation of mouse limbus and choroid following re-analysis of GSE178667. Dashed red line represents the ocular melanocytes based on expression of *Tyr*, and RPE based on expression of *Best1* and *Rpe65.* Feature plot showing mRNA expression of *Tyr, Pax3*, *Ednrb* and *Kit*. (**B**) UMAP representation of Human RPE and choroid samples (GSE135922), visualized using Spectacle (<https://singlecell-eye.org/app/spectacle/>). Violin plots representing expression of *PAX3, FOXD3, SOX10, EDNRB, KIT, TFE3* and *MITF* across cell types, as labeled.

**Figure S21: Immunofluorescent staining for Pax3 in the mouse limbus.** (**A**) Representative 20x image of the mouse eye showing Pax3 positive staining in the limbus region. Sections were counterstained with DAPI to visualize nuclei. (**B**) Post-image processing showing the retina (red), cornea (orange) and limbus (purple), Pax3-positive cells in green. Scale bar = 20μm.
